# Extensive folding variability between homologous chromosomes in mammalian cells

**DOI:** 10.1101/2024.05.08.591087

**Authors:** Ibai Irastorza-Azcarate, Alexander Kukalev, Rieke Kempfer, Christoph J. Thieme, Guido Mastrobuoni, Julia Markowski, Gesa Loof, Thomas M. Sparks, Emily Brookes, Kedar Nath Natarajan, Stephan Sauer, Amanda G. Fisher, Mario Nicodemi, Bing Ren, Roland F. Schwarz, Stefan Kempa, Ana Pombo

**Affiliations:** Max-DelbruDck-Center for Molecular Medicine in the Helmholtz Association (MDC), Berlin Institute for Medical Systems Biology (BIMSB), Epigenetic Regulation and Chromatin Architecture Group, 10115 Berlin, Germany; Humboldt-Universität zu Berlin, Berlin, Germany; Max-Delbrück Centre for Molecular Medicine, Berlin Institute for Medical Systems Biology, Proteomics and Metabolomic Platform, 10115 Berlin, Germany; Max-Delbrück Centre for Molecular Medicine, Berlin Institute for Medical Systems Biology, Evolutionary and Cancer Genomics Group, 10115 Berlin Germany; MRC Laboratory of Medical Sciences, Imperial College London, London W12 0NN, UK; School of Biological Sciences, University of Southampton, Southampton, UK; Dipartimento di Fisica, Università di Napoli “Federico II”, and INFN, Napoli, Italy; Center for Epigenomics and Department of Cellular and Molecular Medicine, University of California, San Diego School of Medicine, La Jolla, CA, USA; Institute for Computational Cancer Biology (ICCB), Center for Integrated Oncology (CIO), Cancer Research Center Cologne Essen (CCCE); BIFOLD – Berlin Institute for the Foundations of Learning and Data, Berlin, Germany

## Abstract

Genetic variation and 3D chromatin structure have major roles in gene regulation. Due to challenges in mapping chromatin conformation with haplotype-specific resolution, the effects of genetic sequence variation on 3D genome structure and gene expression imbalance remain understudied. Here, we applied Genome Architecture Mapping (GAM) to a hybrid mouse embryonic stem cell (mESC) line with high density of single nucleotide polymorphisms (SNPs). GAM resolved haplotype-specific 3D genome structures with high sensitivity, revealing extensive allelic differences in chromatin compartments, topologically associating domains (TADs), long-range enhancer-promoter contacts, and CTCF loops. Architectural differences often coincide with allele-specific differences in gene expression, mediated by Polycomb repression. We show that histone genes are expressed with allelic imbalance in mESCs, are involved in haplotype-specific chromatin contact marked by H3K27me3, and are targets of Polycomb repression through conditional knockouts of Ezh2 or Ring1b. Our work reveals highly distinct 3D folding structures between homologous chromosomes, and highlights their intricate connections with allelic gene expression.

## Introduction

Mammalian cells contain two parental chromosome copies, each with extensive heterozygous sequence variations. Genetic diversity of parental alleles confers advantages over repressive mutations, and is associated with longer lifespan (Xu *et al*, 2019) and reduced risk of aging-related diseases (Belloy *et al*, 2020). Many repressive heterozygous variants, with broad cell functions, are also found in healthy individuals (Schmenger *et al*, 2022), and loss of heterozygosity and single allele amplifications are features of many cancers (LaFramboise *et al*, 2005; Nichols *et al*, 2020). Skewed allelic gene expression has been reported to affect 6% to 80% of genes, depending on the species and tissue (Dixon *et al*, 2015; Crowley *et al*, 2015; Murata *et al*, 2012; Pinter *et al*, 2015; Chen *et al*, 2016; Savol *et al*, 2017; Cleary & Seoighe, 2021). DNA methylation at gene promoters or transcription factor binding sites has been implicated in allele-specific expression of imprinted genes in mouse and human (Noordermeer & Feil, 2020). More recently, genome-wide analyses of mono-allelic expression in the murine zygote, morula and blastocyst, revealed a more prominent role of Polycomb repression than DNA methylation in allelic imbalance of gene expression (Santini *et al*, 2021; Inoue *et al*, 2017). The relative contributions of genetic and epigenetic mechanisms to random allelic expression imbalance are less well understood, but suggested to be highly gene specific (Crowley *et al*, 2015; Marion-Poll *et al*, 2021).

Little is known about haplotype-specific differences in 3D genome structure and their contributions to allelic asymmetries in gene expression. The sparsity of genetic variation between haplotypes makes it technically challenging to map 3D genome structure with haplotype specificity by either sequencing or imaging technologies. In ligation-based methods, such as Hi-C, the unequivocal assignment of ligation events to the correct haplotype (phasing) requires the presence of at least one SNPs on either side of the ligation product and therefore has inherently low sensitivity (reviewed in Li *et al*, 2021). Nevertheless, phased Hi-C data has revealed intrinsic parental variability for the mammalian female X chromosomes, upon random inactivation, and in the timing of chromatin folding during meiosis, but few structural differences have been reported between the two parental copies of somatic chromosomes, except at a small number of imprinted genes (Giorgetti *et al*, 2016; Han *et al*, 2020; Tan *et al*, 2021; Rao *et al*, 2014; He *et al*, 2023; Reinius & Sandberg, 2015; Ferguson-Smith, 2011).

GAM is a ligation-free technology which captures long-range chromatin interactions spanning whole chromosomes and has revealed extensive specificity in the 3D chromatin structure of specific cell types (Beagrie et al, 2017; Beagrie et al, 2023; Winick-Ng et al, 2021; Fiorillo et al, 2021). GAM measures 3D genome topology by sequencing the DNA content from a collection of thin nuclear cryosections, and infers 3D chromatin contacts from the probability of co-segregation of genomic regions across the collection of nuclear slices. As whole genomic regions (typically 20-50 kb long) are called positive in GAM data from the accumulation of many sequencing reads, including many SNP-containing reads, we reasoned that the phasing of GAM data should be highly efficient. Local haplotype fidelity of GAM data has been previously shown (Markowski *et al*, 2021), supporting our efforts to generate haplotype-specific insights into chromatin folding from GAM data.

To investigate differences in the 3D genome folding of homologous chromosomes, we applied GAM to a hybrid mESC line with high SNP density. We developed novel computational pipelines to phase GAM data, and discovered extensive 3D structural differences between the two parental chromosomes across all length scales, including in A/B compartments, topologically associating domains (TADs), and at contact level. We also collected total RNA-seq data and found that 15% of expressed genes have allele-specific expression (ASE) bias in mESCs, including some imprinted genes, but also many housekeeping, ribosomal and histone genes. ASE genes were often located in regions with haplotype-specific structural differences, which coincided with H3K27me3 occupancy, haplotype-specific enhancer-promoter contacts, or CTCF loops. We also inferred chromatin compaction from GAM data, and found that the most active alleles are consistently less condensed than the least active ones. We discovered that many histone genes are ASE genes in mESCs, and that ASE histone genes feature allele-specific long-range chromatin contacts marked by H3K27me3 occupancy. Finally, we performed conditional knockouts of Polycomb enzymatic subunits and showed that the expression of many histone genes is regulated by Polycomb repression mechanisms.

## Results

### Overview of datasets collected

To investigate haplotype-specific differences in 3D genome structure using GAM, we collected data from the F123 mESC line (Gribnau *et al*, 2003). The F123 line was originally derived from F1 hybrid embryos from a CAST/S129 cross and its genotype has high SNP density (average 1 SNP/124 nucleotides across autosomes; **Figure 1a**). GAM data was produced in multiplex mode which combines three independent nuclear profiles (3NP) in each GAM sample (Beagrie et al, 2023; Winick-Ng et al, 2021), and collected from two biological replicates. After quality control, the replicate datasets were merged, resulting in the largest GAM dataset to date, obtained from approximately 3,700 single mESCs (Suppl. Fig. 1a).

**Figure 1:**
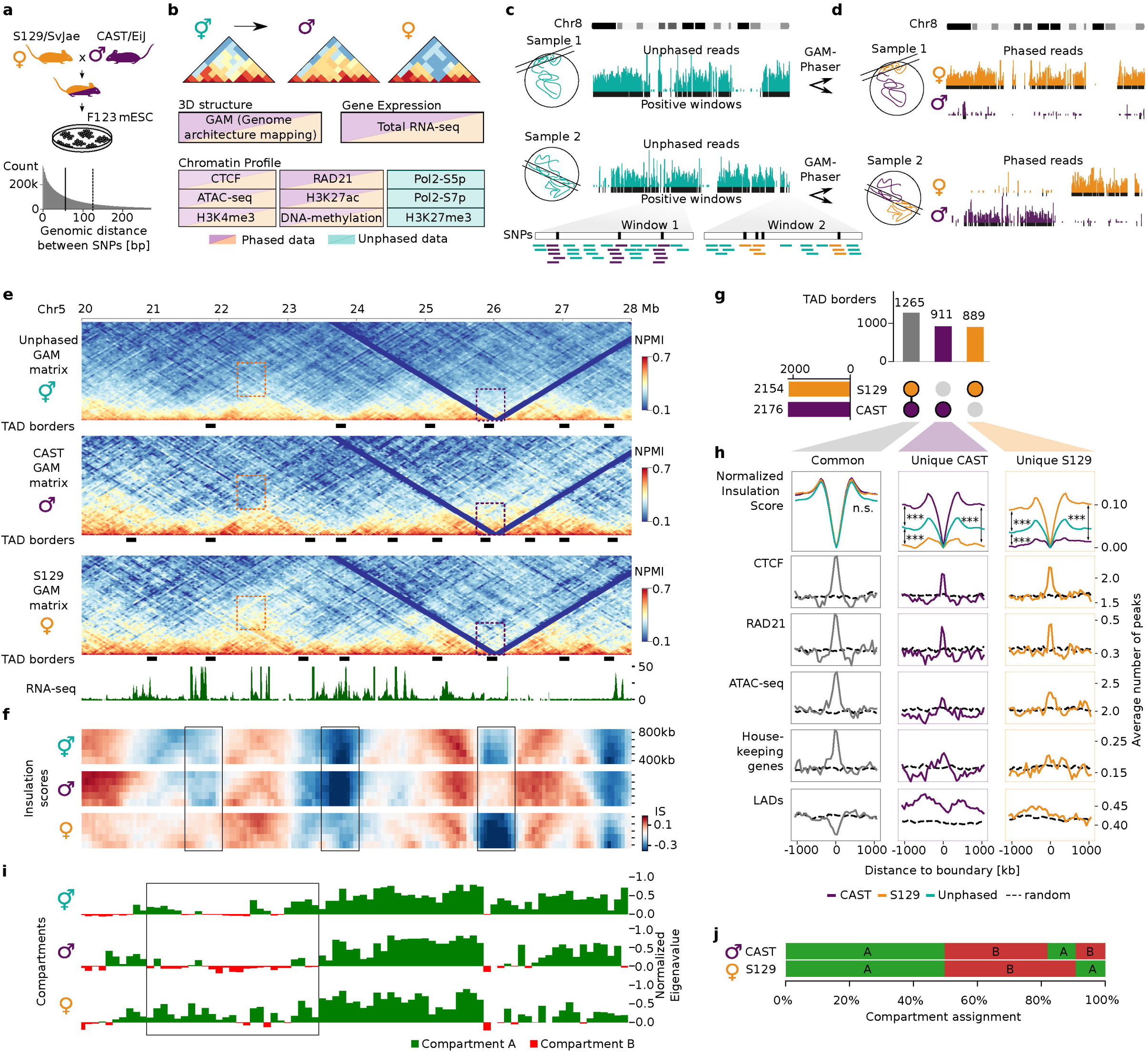
GAM shows structural differences between alleles. **a**, F123 mESC are derived from the cross between S129 and CAST mice. In the F123 genome the median SNP distance is 55 bp (solid line), and the mean is 124 bp (dashed line). **b**, Overview of the data used in this study. GAM, total RNA-seq, ChIP-seq for CTCF, RAD21, H3K27ac and H3K4me3, ATAC-seq and DNA methylation data was collected and phased. Pol2-S5p, Pol2-S7p and H3K27me3 data was collected and analyzed without phasing. **c**, Schematics showing phasing from two GAM samples. Reads mapped to chr8 are shown together with positive windows. Each window can contain a different number of SNPs, and reads mapped to these regions are used for GAM phasing. **d**, Phasing shows that most of the reads belong to one of the haplotypes; black bars below the phased reads represent phased positive genomic windows. Large sections of the chromosome are phased in GAM data. **e**, Unphased and phased GAM maps, TAD borders and the total RNA-seq track are shown for chr5: 20-28 Mb. Colored rectangles mark differences in chromatin contacts between the CAST and S129 haplotypes, with orange and purple corresponding to increased number of contacts for S129 or CAST, respectively. **f**, Heatmap of insulation scores calculated with square sizes that range from 400 to 800 kb. The insulation score heatmaps are represented for the same region as in (**e**). Boxes highlight regions with structural differences between CAST and S129. **g**, UpSet plot shows the number of common and unique TADs to each haplotype. **h**, The normalized insulation score in TADs categorized according to (**g**) is represented in a genomic window centered on the TSS ±1,000 kb). Significance of insulation differences determined with Mann-Witney test; *** represents p < 10^−13^ for all comparisons: common borders, CAST against S129: 0.83; CAST against unphased: 0.41; S129 against unphased: 0.28; CAST unique borders, CAST against S129: 8.9×10^−46^; CAST against unphased: 2.7×10^−14^; S129 against unphased: 1.7×10^−18^; S129 unique borders, CAST against S129: 1.3×10^−47^; CAST against unphased: 1.4×10^−17^; S129 against unphased: 5.6×10^−17^). Other plots show the average number of CTCF, RAD21 and ATAC-seq peaks, Housekeeping genes and LADs around the TSS. Dashed lines depict the expected number of features using circular permutations and averaging the score from 10 iterations. **i**, A and B compartment annotations and normalized eigenvector values for unphased and phased matrices for the region shown in panel **e**. Box highlights a region with notable differences between CAST and S129 matrices. **j**, Compartment assignments show 18% different annotations in CAST and S129.

To address the impact of haplotype-specific 3D genome structure on gene expression and chromatin regulation, we also mapped gene expression using total RNA-seq, chromatin occupancy using ChIP-seq of RNA polymerase II phosphorylated on Serine-5 (Pol2-S5p) or Serine-7 (Pol2-S7p) residues of its C-terminal domain, and the Polycomb mark H3K27me3 (**Figure 1b**). We collected published datasets produced in F123 mESCs for ChIP-seq of CTCF, cohesin (RAD21), H3K4me3 and H3K27ac (Huang *et al*, 2021), chromatin accessibility (ATAC-seq; Juric *et al*, 2019), and DNA methylation (WGBS; Li *et al*, 2019). Finally, we also considered published annotations of lamina-associated domains (LADs) obtained by Lamin B1 DamID from the mESCs clone E14Tg2A (Peric-Hupkes *et al*, 2010). The datasets produced, publicly available and the processed data resources are summarized in SI Table 1.

### GAM-phaser: a pipeline to phase GAM data

We developed the GAM-Phaser pipeline to phase GAM data to the CAST and S129 haplotypes (Suppl. Fig. 1b). Briefly, the positive genomic windows in each GAM sample are first defined in unphased GAM datasets (**Figure 1c**), using a threshold of nucleotide coverage as previously described (Winick-Ng *et al*, 2021). Next, SNP-containing reads are phased to CAST and S129 haplotypes and the same threshold is applied to call haplotype-specific windows in each GAM sample (**Figure 1d**). After phasing genomic windows, it becomes possible to distinguish whether a given GAM sample contains DNA from one or both chromosome copies (Sample 1 and 2, respectively; **Figure 1d)**

The overall read-phasing efficiency achieved across all F123 GAM sequencing datasets was 37% to CAST or S129 haplotypes (Suppl. Fig. 1c). However, it was possible to phase 70 to 75% genomic windows at 10 to 100 kb resolutions, respectively, evenly detected between CAST and S129 windows (Suppl. Fig. 1d). The high efficiency of GAM data phasing results from the presence of multiple nucleotide polymorphisms, which collect many SNP-containing reads, in each positive genomic window. In comparison, phasing of Hi-C data from equivalent hybrid mESC lines has achieved only 26 or 35% phasing efficiency of ligation events (Giorgetti *et al*, 2016; Glusman *et al*, 2014; Bonora *et al*, 2021), which we confirm in published Hi-C data from a hybrid mESC line with similar SNPs density (26%; Suppl. Fig. 1e). Genomic windows were rarely phased to both CAST and S129 haplotypes in the same nuclear profile (Suppl. Fig. 1d). To exemplify the low expected co-detection of allelic windows in the same nuclear slices, we took published imaging data using fluorescence in situ hybridization in cryosections (cryoFISH) of a different ESC clone (46C) targeting the genomic regions containing *Hoxb1* or *Hoxb13* with fosmid probes covering approximately 40 kb (Barbieri *et al*, 2017; SI Table 2). CryoFISH data analyses confirmed that a minority of nuclear sections contain both copies of each locus (13% and 11%, respectively; Suppl. Fig. 1f).

To determine a suitable resolution of the phased GAM data, we calculated the detectability of window co-segregation events at different genomic resolutions (Suppl. Fig. 1g). We chose a window resolution of 50 kb for downstream analyses, which gives >97% detection of co-segregation events across all genomic distances. Higher resolutions down to 10 kb also gave good co-segregation frequencies for unphased data; for example, 99.28% of all possible pairs of 10 kb genomic windows within 10 Mb were co-detected in at least one GAM sample.

### GAM detects extensive haplotype-specific differences in chromatin contacts

To begin assessing the extent of haplotype-specific differences in chromatin contacts captured in GAM data, we compared unphased and phased contact matrices (**Figure 1e**). We found extensive structural variability between the CAST– and S129-phased matrices, and noticed that both local and long-range contacts are stronger and more obvious in the haplotype-specific matrices than in the unphased, average matrices (**Figure 1e**, orange and purple rectangles, respectively for strong contacts in S129 or CAST haplotypes). Structural variability between haplotypes becomes even more prominent when plotting whole chromosome matrices, where clusters of increased long-range contacts are clearly visible across genomic distances (Suppl. Fig. 2a, orange and purple arrows, respectively). Contact distance decay and momentum curves showed similar frequency of contacts between haplotypes within < 5Mb of genomic distance, but became visibly distinct at long-range distances with different haplotype preferences depending on the chromosome (Suppl. Fig. 2b).

### Most TAD borders are haplotype specific

To quantify haplotype-specific differences at the level of TAD organization, we calculated insulation scores at different length scales, using 400 to 800 kb square sizes (Crane *et al*, 2015; Winick-Ng *et al*, 2021), and found clear differences in insulation between parental genomes (**Figure 1f**; see boxes and Suppl. Fig. 2c for an additional example; for insulation score data see permanent data repository Irastorza-Azcarate *et al*., 2024). Consistent with the unphased matrices being an average of the CAST– and S129-specific contact matrices, we confirmed that the CAST and S129 insulation scores correlated less with each other than with unphased insulation scores (400 kb insulation square sizes; Suppl. Fig. 2d).

Next, we computed TAD borders in unphased and phased matrices using the 400 kb insulation square size, as previously described (Winick-Ng *et al*, 2021). More than 40% of all TAD borders detected are haplotype specific (911 and 889 unique to CAST and S129) compared with 1,265 common borders detected in both haplotypes (**Figure 1g**, SI Table 3; for all combinations see Suppl. Fig. 2e). The distinct insulation between haplotype-specific TAD borders was confirmed by comparing average insulation plots (Mann-Witney test, *** p < 10^−13^ for all comparisons; **Figure 1h**). Many borders common to both haplotypes were also detected in unphased matrices (1061), as expected, and some CAST– and S129-specific borders could also be detected in the unphased matrices but not in the other haplotype (373 and 351, respectively), likely reflecting their strong prevalence in one haplotype chromosome across the cell population (Suppl. Fig. 2e). However, many CAST– and S129-specific borders were not captured in the unphased matrices (514 and 518, respectively), highlighting the specificity of chromatin topology in the two haplotypes.

We asked whether haplotype-specific TAD borders were enriched for CTCF, cohesin or housekeeping (HK) genes, as previously shown for unphased TAD borders (Dixon *et al*, 2012). CTCF and cohesin were found highly enriched in both common and haplotype-specific borders, whereas housekeeping genes and chromatin accessibility (ATAC-seq) are more strongly enriched in common borders. We also noted a preference for common borders to more likely correspond to LAD/interLAD transitions (22.9%) than CAST– and S129-unique borders (14.4% or 11.4%, respectively; Suppl. Fig. 2f).

### Haplotype-specific compartments account for 20% of the genome

Previous work in mouse T cells from B6×CAST hybrid mice detected 4% of compartment changes between haplotypes (Han *et al*, 2020), and region-specific examples of compartment changes have also been reported at specific loci in hybrid mESCs (Rivera-Mulia *et al*, 2018). To quantify the extent of haplotype differences in compartment A/B annotation genome-wide, we computed eigenvector values from principal component analysis (PCA) from unphased and haplotype-specific GAM matrices (**Figure 1i**, box; see also whole chromosome regions in Suppl. Fig. 2g; SI Table 4). Genome-wide analyses showed that most genomic regions have the same compartment definition in CAST and S129 (49% and 32% of genomic regions being annotated A-A or B-B, respectively), with 18% varying between CAST and S129 (9% for A-B, and vice versa; **Figure 1j**). We observed that the distributions of eigenvector values of allele-specific matrices are more symmetrical compared to unphased values and cover a wider range, suggesting that the phased data captures compartmentalization states better than the haplotype-averaged unphased data (Suppl. Fig. 2h).

To investigate the functional consequences of extensive haplotype-specific differences in chromatin structure on gene expression and their relationship with chromatin-based mechanisms of gene regulation, we quantified the haplotype-specific differences in gene expression and chromatin features, and characterized their co-occurrence across the linear genome. Subsequently, we integrated allele-specific 3D genome structure with the linear distribution of allele-specific gene expression and chromatin occupancy.

### Allele-specific expressed genes are enriched in housekeeping, ribosomal and histone gene groups

To understand the extent of allele-specific gene expression in F123 mESCs, we measured gene expression from total RNA-seq data for protein-coding and long non-coding genes, after selecting the most expressed transcript isoform (based on the levels of Pol2-S5p and Pol2-S7p at annotated transcription start sites; see Methods). We calculated differential allelic expression as previously described (Castel *et al*, 2015), considering both exonic and intronic regions (for gene expression levels see Irastorza-Azcarate *et al*., 2024). Out of 17,956 expressed genes, we detected 13,713 genes similarly expressed from both alleles, 2,222 genes with ASE imbalance (|log2 fold change| ≥1, adjusted p-value <0.05), of which 1,308 and 914 were more expressed from the CAST or S129 genomes, respectively, and 2,021 expressed genes without SNP (**Figure 2a**). Among the ASE genes, we found 193 monoallelic genes, including several histone genes, such as *Hist2h2ac* expressed from CAST paternal haplotype (**Figure 2b**), and 15 imprinted genes, including *Lin28a* and *Peg13*, more expressed from the CAST allele, and *Cdkn1c*, more expressed from the S129 allele. ASE genes more frequently exhibited higher expression from the paternal allele, as previously reported in murine embryonic fibroblasts and adult tissues (Crowley *et al*, 2015; Savol *et al*, 2017).

**Figure 2:**
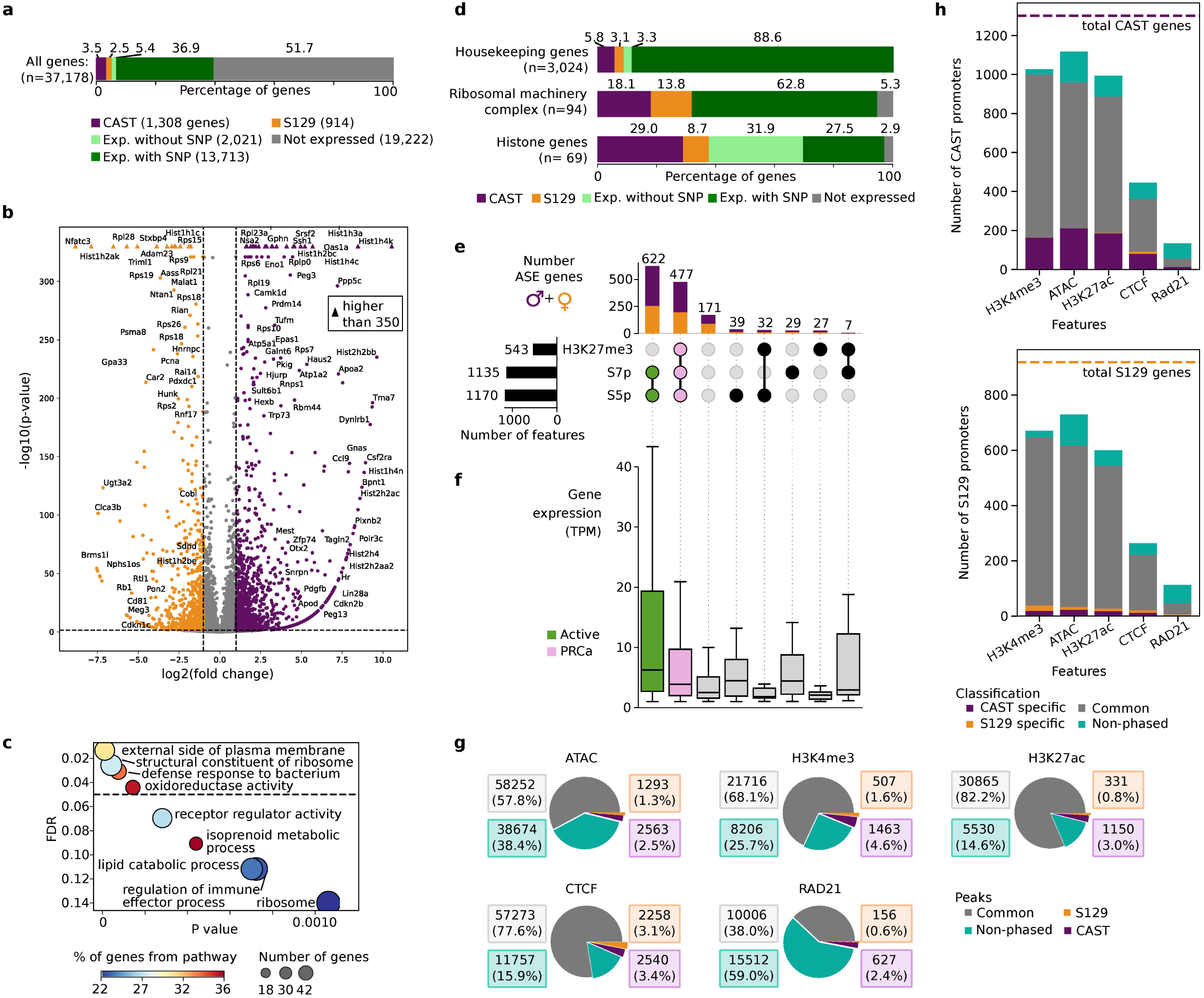
Allele-specific expressed (ASE) genes are enriched for housekeeping, ribosomal protein and histone genes, and many contain Polycomb. **a**, Number and percentage of CAST and S129 ASE genes expressed with no SNP, genes that are biallelic and genes that are not expressed. **b**, Volcano plot of all expressed genes containing SNPs. Genes with a |log2 fold change| ≥1 and an adjusted p-value of <0.05 were classified as ASE. **c**, Significant Gene Ontology (GO) terms of ASE genes. **d**, Bar plot showing percentage of genes belonging to housekeeping, ribosomal machinery complex and histone protein gene groups. **e**, Overlap of CAST and S129 ASE genes with H3K27me3, Pol2-S5p and Pol2-S7p. **f**, Gene expression of the different groups in panel (**e**) using TPM. **g**, Number and percentage of H3K4me3, H3K27ac, CTCF and RAD21 ChIP-seq and ATAC-seq peaks that are CAST or S129 specific, common or could not be phased. **h**, Number of CAST (top) and S129 gene promoters (bottom) that overlap with different features.

Gene Ontology (GO) enrichment analysis showed that ASE genes are involved in metabolic processes, immunity response and the ribosomal machinery (**Figure 2c**; SI Table 5). Separate GO enrichment analysis of CAST and S129 ASE genes showed no haplotype-specific enrichment of biological functions. Amongst the ASE genes, we found many ribosomal protein genes (32%, 30/94; **Figure 2d**), a group of genes previously reported to be expressed with allelic imbalance in other mouse tissues or cell types, and in Medaka and catfish tissues (Murata *et al*, 2012; Pinter *et al*, 2015; Crowley *et al*, 2015; Chen *et al*, 2016). The ASE list also contained 38% of all histone genes (26/69) and 8% of housekeeping genes (SI Table 6). The ASE imbalance of histone genes has been reported in mouse embryonic fibroblasts, and can also be observed by mining publicly available resources from mouse tissues (Crowley *et al*, 2015; Pinter *et al*, 2015; Savol *et al*, 2017), but has so far not been investigated.

### H3K27me3 occupies a third of ASE gene promoters

ASE imbalance is thought to be achieved by repression mechanisms acting on one allele (Garcia *et al*, 2014), and some studies report a major role of Polycomb repression in monoallelic expression, predominantly at the maternal allele (Inoue *et al*, 2017; Santini *et al*, 2021). To explore whether Polycomb repression mechanisms are also important at ASE genes, we mapped H3K27me3, Pol2-S5p and Pol2-S7p occupancy. Regions enriched for these markers cover extended genomic stretches (median length 2,000-3,400 bp; Suppl. Fig. 3a), with few peaks containing only CAST or S129 reads (0.6-0.8 % and 0.3 % for CAST and S129 peaks, respectively; for full list see Irastorza-Azcarate *et al*., 2024), as shown previously (Savol *et al*, 2017). We therefore chose to explore the contribution of Polycomb and Pol2 to ASE imbalance using unphased chromatin occupancy and classifying the promoters of ASE genes according to their presence or absence.

Previous genome-wide analyses in mESCs showed that the promoters of signaling or metabolic genes are often co-occupied by H3K27me3, Pol2-S5p and Pol2-S2p, reflecting a mixed Polycomb-Active (PRCa) promoter state suggested to result from different promoter states between alleles or between cells (Brookes *et al*, 2012; Ferrai *et al*, 2017). We classified all non-overlapping gene promoters in F123 mESCs according to their H3K27me3, Pol2-S5p or Pol2-S7p occupancy (see Irastorza-Azcarate *et al*., 2024), and found that one third of ASE genes have PRCa promoter states (H3K27me3+S5p+S7p+, 477 genes; **Figure 2e**), including signaling and metabolic genes, such as *Mapk13* and *Apoe*, respectively. The expression of ASEs marked by H3K27me3, S5p and S7p is approximately half of the expression levels of ASE genes occupied by S5p and S7p only (**Figure 2f**), consistent with repressive effects of Polycomb acting on one allele. These observations suggest a major role for Polycomb repression mechanisms in ASE imbalance at a large subset of ASE genes, potentially to dynamically regulate signaling or metabolic responses.

### Active chromatin features are mostly biallelic and not associated with allelic imbalance

To further investigate other chromatin-mediated mechanisms that might contribute to haplotype-specific chromatin structure, we mapped chromatin accessibility (ATAC-seq) and H3K4me3, H3K27ac, CTCF and cohesin (RAD21) occupancy (see Irastorza-Azcarate *et al*., 2024). Occupancy peaks were relatively short (305-1,507 on average (Suppl. Fig. 3a), and most could be phased (41-86%), but were biallelic (**Figure 2g**). A minority of peaks were classified as CAST (2.5-3.8%) or S129 (0.6-3.1%) specific, and were slightly more abundant in the CAST haplotype, a preference also observed in the number of CAST ASE genes.

We measured the overlap between ASE gene promoters and haplotype-specific ATAC, H3K4me3, H3K27ac, CTCF or RAD21 peaks (**Figure 2h**). Most ASE gene promoters coincide with biallelic peaks of ATAC, H3K4me3 and H3K27ac, less frequently with CTCF, and rarely with RAD21. CAST-specific peaks that overlap with ASE genes are preferentially found at promoters of genes more highly expressed from the CAST haplotype (1-16% depending on chromatin feature), in contrast with S129-specific peaks which rarely coincide with S129-specific promoters (0.4-2%). This tenfold haplotype imbalance is unlikely to be technical, as for example the detection of CAST and S129 ATAC peaks is almost even (2.5% and 1.3%, respectively). We also considered ATAC peaks outside of gene promoters but in proximity to ASE genes, and found that CAST ATAC peaks are preferentially nearest to CAST ASE genes and S129 ATAC peaks closer to S129 ASE genes (Suppl. Fig. 3b). The observation that both CAST– and S129-specific ATAC peaks have a preference for proximity to ASE genes more expressed in the same haplotype suggests a role for enhancer-promoter (E-P) chromatin contacts in ASE imbalance.

We searched for transcription factor motif enrichment at CAST or S129 ATAC peaks present at promoters, intergenic or genic regions, and found a single transcription factor, ZFP57, enriched in CAST-specific peaks at CAST gene promoters, and not in other promoters or genomic regions (Suppl. Fig. 3c; for list of motifs in ATAC-seq peaks see Irastorza-Azcarate *et al*., 2024). ZFP57 is a zinc finger protein involved in maintenance of imprinted genes through binding of DNA methylated regions (Mackay *et al*, 2008, Shi *et al*, 2019). To explore the association of ASE imbalance with differential methylation in F123 mESCs, we called differentially methylated genes from published phased whole bisulfite sequencing data in F123 mESCs (Li *et al*, 2019). Only 61 ASE gene promoters were found associated with allele-specific DNA methylation (2.7% of all ASE genes; Suppl. Fig. 3d, SI Table 7), including three imprinted genes, *Mest, Snrpn* and *Peg13* (SI Table 6). *Mest* is a CAST ASE gene which shows stronger chromatin contacts at the maternal than paternal *Mest* locus (Suppl. Fig. 3e), in line with previous reports in neonatal and adult neurons, using Dip-C (Tan *et al*, 2021).

To complete our exploration of linear chromatin features and their association with ASE imbalance, we considered CTCF and RAD21 peaks. Most CTCF peaks are biallelic (57,273), and only 2,540 and 2,258 are CAST or S129 specific, respectively (**Figure 2g**). In contrast, most Rad21 peaks could not be phased (15,512), and only 627 or 156 peaks were assigned to the CAST or S129 alleles, respectively. Although CTCF peaks rarely overlap ASE gene promoters (**Figure 2h**), some allele-specific CTCF peaks overlap promoters of monoallelic expressed genes, for example *Cdkn2b* and *Hist2h4* (for full list of genes and overlapping features see Irastorza-Azcarate *et al*., 2024). These results suggest that CTCF and RAD21 are not a major feature of ASE imbalance. We also noticed that haplotype-specific CTCF peaks are present at approximately one quarter of TAD borders, but they always co-occur with biallelic CTCF peaks, and show no preference for borders of the matching haplotype (Suppl. Fig. 3f,g), suggesting that CTCF-mediated mechanisms are not general drivers of haplotype-specific TAD formation.

### CAST-specific ASE genes and chromatin features co-occur in the linear genome

Previous studies in mouse and medaka tissues reported ASE gene clustering in the linear genome (Garcia *et al*, 2014; Crowley *et al*, 2015). We inspected the position of ASE genes in F123 mESCs across whole chromosomes and confirmed a tendency for ASE gene clustering in mESCs (**Figure 3a**, Suppl. Fig. 4a). Genome-wide analyses showed that CAST and S129 genes are present in all autosomal chromosomes, tend to be clustered, and are often intermingled with each other (circular Permutation test, p-values = 0.0001, 0.0145, 0.0001 for CAST, S129 and CAST+S129; Suppl. Fig. 4b). ASE genes are located in genomic regions with high density of expressed genes compared with regions without ASE genes (t test: p-value = 2.7×10^−71^; Suppl. Fig. 4c).

**Figure 3:**
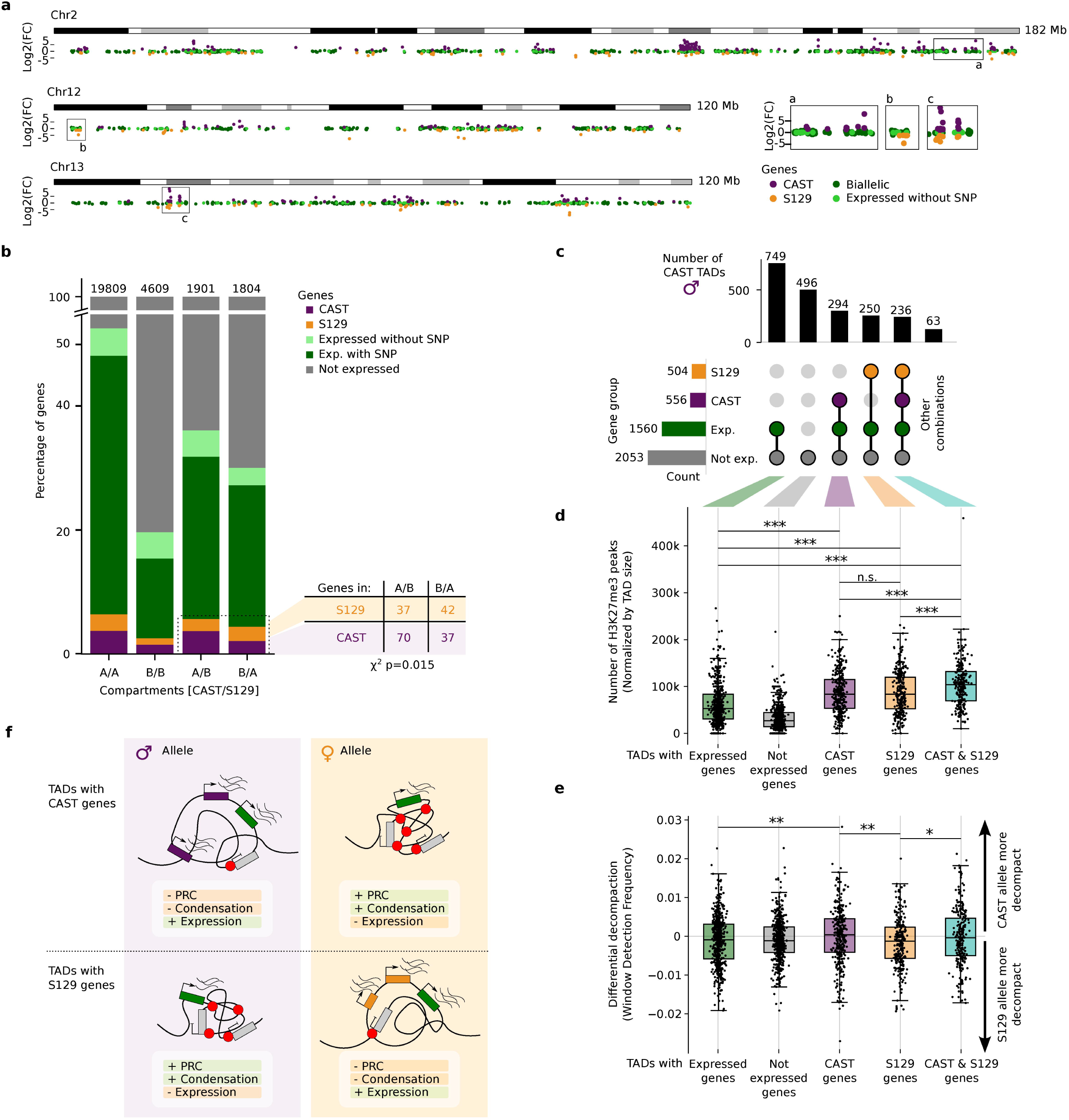
ASE genes are clustered in TADs enriched for H3K27me3 occupancy. **a**, Manhattan plot showing log2 fold change of all expressed genes for chromosomes 2, 12 and 13. Genes are colored according to their expression as CAST or S129 specific, biallelic or do not contain SNPs. Boxes show regions that were zoomed: a region with majority of CAST ASE genes, majority of S129 ASE genes or a mix of CAST and S129 ASE genes. **b**, Bar plot showing the percentage of genes that overlap with A or B compartments in both haplotype, or have different compartment annotations in CAST and S129. The preferred tendency for CAST genes to be in CAST compartment A, and S129 genes to be in S129 compartment A is statistically significant (Chi-square test= 0.015). **c**, UpSet plots showing for the CAST allele (S129 allele in sup fig 5c), groups of TADs containing different sets of types of genes and their number. **d**, For each group in panel c, the number of H3K27me3 peaks normalized by TAD size. (two-sided t-test: *p<0.05, **p<0.01, ***p<0.001; p-values from top to bottom in CAST TADs: 1.0×10^−16^, 7.0×10^−15^, 1.9×10^−32^, 4.4×10^−5^, 7.6×10^−5^, n.s: 0.95. **e**, The differential (CAST-S129) window detection frequency is represented for each group in panel (**c**). Negative values indicate decompaction in the S129 haplotype, while positive values indicate decompaction in CAST (two-sided t-test: *p<0.05, **p<0.01, ***p<0.001; p-values from top to bottom for CAST TADs: 0.008, 0.003, 0.018. **f**, Summary model displaying differences in chromatin compaction of TADs containing CAST or S129 ASE genes.

Next, we measured the genomic overlap and clustering of ASE genes and active chromatin features, and found that CAST genes and chromatin features often co-occur with each other, in contrast with S129 features and genes which rarely co-occur (Suppl. Fig. 4d). CAST features and CAST genes do not frequently co-occur with S129 features or S129 genes, indicating that CAST and S129 features are generally segregated along the linear genome (Suppl. Fig. 4e).

Taken together, our exploration of the linear organization of ASE genes and chromatin accessibility and occupancy suggest that different mechanisms may control ASE gene expression, of which Polycomb repression was most prevalently associated with ASE imbalance. In the next sections, we investigated how these linear genome features relate with haplotype differences in 3D genome structure.

### ASE clustering occurs preferentially within compartment A

We asked whether ASE gene clustering was reflected in haplotype-specific compartment transitions. ASE genes are most abundantly present within compartment A (euchromatic) annotations which are common to both haplotypes (Suppl. Fig. 5a, circular Permutation test, 10,000 permutations, p-value = 0.0001), a tendency that is likely driven by their preferred co-occurrence with biallelic expressed genes. However, 107 CAST and 79 S129 ASE genes are present in genomic regions with A-B or B-A (CAST-S129) compartment assignments, with a preference for CAST and S129 genes being present in compartment A annotations of the corresponding haplotype (**Figure 3b**; Chi-squared test, p = 0.01125). Haplotype-specific ATAC, CTCF, H3K4me3 and H3K27ac peaks were also found preferentially associated with haplotype-specific compartment differences, with CAST-specific peaks being slightly more abundant in A-B (CAST-S129) compartments, and S129-specific peaks in B-A regions, with the exception of S129-specific CTCF peaks which are equally distributed in A-B and B-A regions (Suppl. Fig. 5b). Although the majority of ASE genes and haplotype-specific chromatin occupancy features are preferentially present in compartment A (euchromatic) regions common to CAST and S129, exceptions can be found that highlight a role of context-based mechanisms in ASE regulation.

### ASE genes are clustered within TADs enriched for H3K27me3 occupancy

We then asked whether ASE gene clustering in the linear genome reflects the TAD organization. ASE genes are present in 45% of TADs, in all cases together with biallelically expressed genes (**Figure 3c**). Approximately one third of TADs contain CAST ASE genes, another third S129 ASE genes, and the last third containing both (**Figure 3c** for CAST TAD annotations, Suppl. Fig 5c for S129 TAD annotations). ASE gene clustering within TADs is statistically significant for CAST, S129 or combined CAST/S129 ASE genes (Permutation test, 10,000 permutations, p-values = 0.0001, 0.0001, 0.0001 respectively; Suppl. Fig. 5d, 5e).

We asked whether TADs containing ASE genes are also enriched for H3K27me3 peaks, and found a statistically significant enrichment compared with TADs containing only biallelic expressed genes or silent genes (t-test, p-values < 0.001; **Figure 3d**, Suppl. Fig. 5f). The H3K27me3 enrichment is especially strong in TAD annotations containing both CAST and S129 ASE genes, compared with TADs containing ASE genes of only one haplotype, suggesting allele-specific local contributions of H3K27me3 to ASE imbalance. Since chromatin compaction is a feature of Polycomb activity *in vitro* and *in vivo* across short and long genomic regions (Nichols *et al*, 2020; Barbieri *et al*, 2017; Schoenfelder *et al*, 2015), we first asked whether the differential presence of CAST or S129 ASE genes within specific TADs correlated with increased chromatin decompaction in the haplotype with the larger number of expressed genes. We measured the frequency of detection of genomic windows, a unique feature of GAM data that infers chromatin condensation based on the increased frequency of detection of decondensed genomic regions (with larger volume) compared with more condensed regions (with lower volume), across the full collection of GAM samples (Beagrie *et al*, 2017; SI Table 8). We found that the average window detection frequency (WDF) of genomic regions within TADs containing only CAST or only S129 ASE genes is on average higher in the most expressed allele irrespective of haplotype (**Figure 3e**, Suppl. Fig. 5g). Increased WDF is also observed at the gene level, as genomic windows containing the most expressed allele are also more decompacted (Fisher’s exact test p = 5.3×10^−5^; Suppl. Fig. 5h). The observation that TADs with ASE imbalance are associated with Polycomb occupancy and compaction of the repressed allele, provide orthogonal support for a role of Polycomb repression in chromatin condensation genome-wide, which we show here in the context of haplotype-specific chromatin regulation (**Figure 3f**).

### Long-range interactions in the Hist1 gene cluster are allele specific

To further explore how the linear clustering of ASE genes relates to allelic differences in higher-order chromatin contacts, we considered the Hist1 locus which contains 19 ASE histone genes. The Hist1 locus is the largest and most dense of the four histone loci, and contains three Hist1 sub clusters (∼200, ∼10 and ∼500 kb) in a 2 Mb region, harboring a total of 55 histone genes interspersed with two silent clusters of sensory receptor genes, Olfr and Vmnr (**Figure 4a**). The Vmnr cluster is annotated as B compartment in unphased GAM data, and a LAD region, flanked by active histone genes in compartment A. Most histone genes in the Hist1 locus are expressed in F123 mESCs (52 out of 55 genes) of which 35 contain SNPs. Of the 19 ASE genes in the locus, 14 and 5 genes are more highly expressed from the CAST or S129 allele, respectively, indicating that the Hist1 cluster is more transcriptionally active in the CAST than the S129 chromosome copy (Suppl. Fig. 6a).

**Figure 4:**
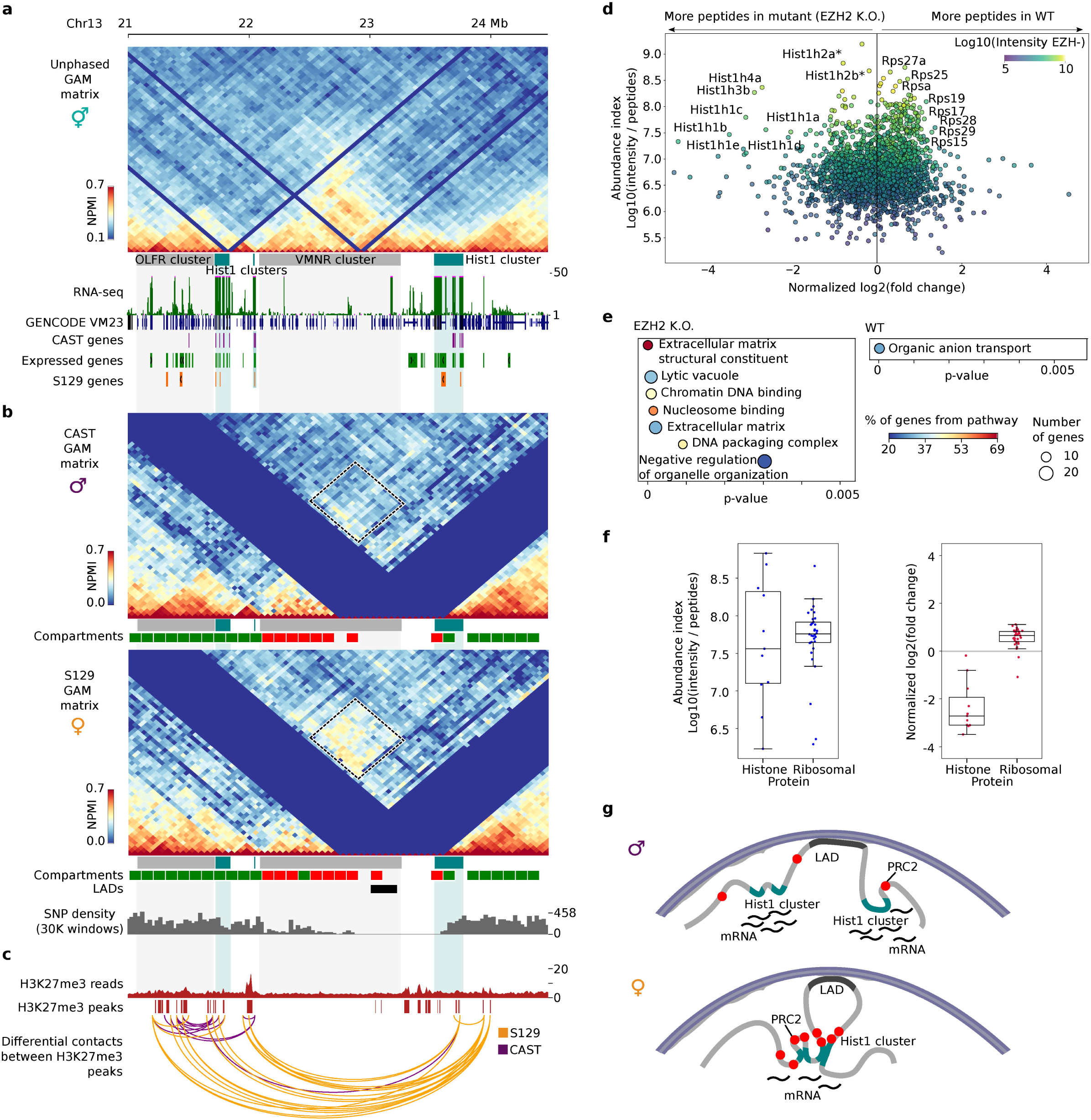
Histone genes in the Hist1 cluster establish S129-specific contacts that coincide with H3K27me3 occupancy, and are regulated by Polycomb repression mechanisms. **a**, Unphased GAM map of the Hist1 locus (chr13: 21-24.5Mb). Below, tracks showing the position of each Hist1 cluster, olfactory receptor cluster and the VMNR cluster; total RNA-seq data, position of all genes, expressed genes and genes specific to CAST and S129 alleles. **b**, Phased maps of the same region to the CAST and S129 allele. Below, SNP density track at 30 kb bins, showing a region which contains part of the VMNR cluster and the rightmost Hist1 cluster, devoid of SNPs. The rectangle highlights contacts between the Hist1 clusters which are strong in the S129 allele, and weak in the CAST allele. **c**, Tracks for H3K27me3 reads and peaks. Below, allele-specific contacts for each allele extracted from the phased GAM maps that coincide with H3k27me3 peaks. **d**, SILAC experiments carried out in ESC-Ezh2-1.3 cells grown in the absence or presence of tamoxifen to induce conditional knockout of *Ezh2* show that histone protein genes are upregulated upon Ezh2 knockout. *Abundance* was calculated as intensity divided by number of peptides, while *normalized log2fc* was calculated applying the z score normalization to the log2 of heavy/light (H/L) ratio of the WT experiment divided by the H/L ratio of the conditional knockout. **e**, Gene Ontology terms for the top 5% upregulated genes for each condition. **f**, Boxplots showing the abundance index and the log2 fold change for histone proteins and ribosomal proteins related to panel d. **g**, Proposed model for the Hist1 locus folding and gene regulation. Haplotype-resolved GAM data shows that the Hist1 clusters come together preferentially in the S129 allele. These contacts may be mediated by Polycomb which establishes a repressive environment and thus results in lower overall expression. The Hist1 clusters in the CAST allele are spatially separated which results in increased gene expression.

Unphased GAM data shows that the three Hist1 locus subclusters interact with each other (**Figure 4a**), establishing long-range contacts that resemble those found in human ESCs by SPRITE (Quinodoz *et al*, 2018). As the Hist1 locus has a robust density of SNPs, except across the *Vmnr* gene cluster, it was possible to phase most of the region. In the haplotype-specific GAM contact matrices, we found that the Hist1 locus shows extensive structural differences between CAST and S129 haplotypes (**Figure 4b**), in particular a large S129-specific patch of strong contacts between the most distant Hist1 subclusters, separated by 1.5 Mb. As the S129 locus expresses fewer genes than the CAST locus, we hypothesized that the long-range contacts might relate to histone gene repression. H3K27me3 occupancy was detected at 11 out of 19 histone ASE genes, and their promoters are classified as PRCa (Suppl. Fig. 6b; see Irastorza-Azcarate *et al*., 2024). Although histone genes have not previously been reported as targets of Polycomb repression mechanisms, evidence for the presence of H3K27me3 or mono-ubiquitinylated H2A (H2Aub1) at histone gene promoters can be traced in published mESC datasets for *Hist3h2ba (*Brookes *et al,* 2012), and *Hist2h3c1*, *Hist2h4*, *Hist3h2ba* genes (Ferrai *et al*, 2017) in different mESC lines.

### Histone genes are upregulated upon conditional Polycomb knockout

To explore potential roles of Polycomb repression in Hist1 gene regulation, we calculated differential contacts between CAST and S129 matrices (Suppl. Fig. 6c), and extracted all allele-specific contacts in the region involving windows containing H3K27me3 peaks (**Figure 4c**, Suppl. Fig. 6d). We found that these long-range contacts connect all 3 clusters in the S129 allele, suggesting that the repression of a larger number of histone genes in the S129 haplotype may relate to local and long-range effects of Polycomb repression.

To directly address a functional role for Polycomb repression in the dampening of histone gene expression, we took advantage of two previously characterized conditional tamoxifen-inducible knockout cells of *Ring1b* (murine ESC-ERT2 clone; Stock *et al*, 2007) or *Ezh2* (murine ESC-Ezh2-1.3 clone; Pereira *et al*, 2010), which encode the major enzymatic activities of Polycomb repressor complex 1 (PRC1) or 2 (PRC2), respectively. Upon addition of tamoxifen, ESC-ERT2 and ESC-Ezh2-1.3 lose H2Aub1 or H3K27me3, respectively, within 24/48 or 96h (Stock *et al*, 2007; Pereira *et al*, 2010). We performed quantitative SILAC mass spectrometry analysis in the two cell lines, before and after knockout induction (SI Table 9). We discovered that histone proteins were highly upregulated after knockout of either Ring1b or Ezh2, and were among the proteins with highest fold change upregulation (**Figure 4d**, Suppl. Fig. 6e). In fact, GO enrichment analysis on proteins with 5% highest fold change shows enrichment for terms associated with DNA packing complex and nucleosome binding proteins (**Figure 4e**, SI Table 5). Finally, we were curious to see whether ribosomal proteins were also affected by the conditional Polycomb knockout. Interestingly, there was not much difference in ribosomal protein expression before or after knockout induction, supporting the view that Polycomb repression has a specific role downregulating histone expression (**Figure 4f**, Suppl. Fig. 6f).

Our observations show that many histone genes are ASE genes regulated by Polycomb repression mechanisms, with promoters occupied by Pol2-S5p, –S7p and H3K27me3. We also show that histone genes within the Hist1 locus establish long-range chromatin contacts, often occupied by H3K27me3, which bridge a gene-silent LAD, and occur especially in the S129 haplotype that expresses fewer Hist1 genes (see schematics in **Figure 4g**).

### Allele-specific contacts between ASEs and enhancers

Finally, we were curious about allele-specific contacts between ASE gene promoters and putative regulatory regions (enhancers; E), and whether E-ASE gene contacts would be predominant in the most or least expressed allele. To define a stringent list of E-ASE gene contacts, we selected the strongest contacts (z-scores higher than 2.0; Suppl. Fig. 7a) that connect ASE genes within 2Mb, and which were also marked by Pol2-S5p and ATAC peaks, with putative enhancers that were marked by Pol2-S5p, ATAC and H3K27ac (**Figure 5a**; the table of differential contacts and features is available in GSE254717).

**Figure 5:**
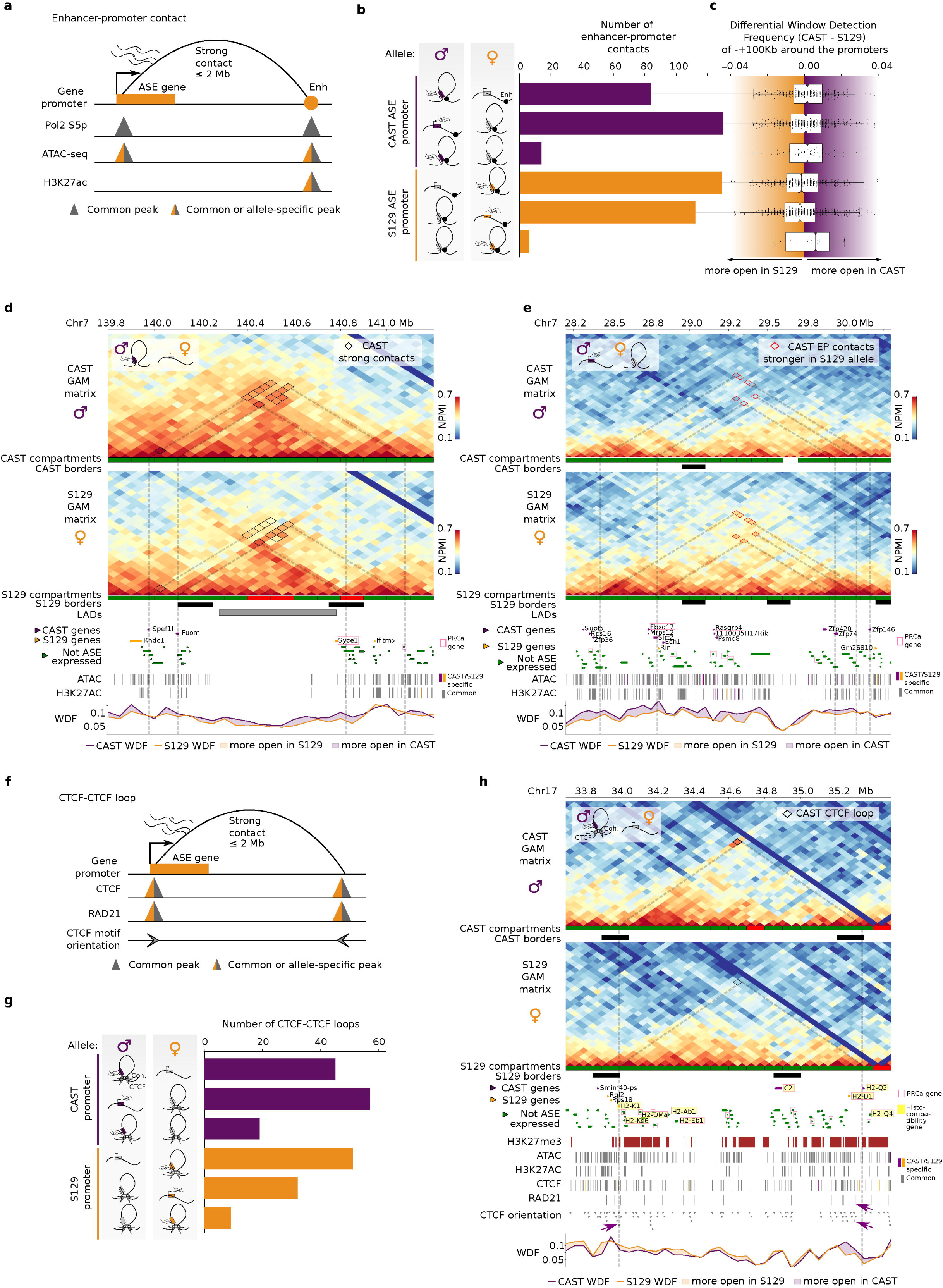
Enhancer-promoter contacts and CTCF-loops coincide with ASE genes. **a**, Features and conditions used to define enhancer-promoter contacts (Enh, Enhancer). **b**, Quantification of enhancer-promoter contacts, depending on the different configurations in each allele. **c**, Boxplot showing the normalized window detection frequency (WDF) for each configuration. **d**, Example region with contact differences on chr7 with allele-specific enhancer-promoter contacts. **e**, Example for decondensation in the CAST allele with allele-specific enhancer-promoters (EP) contacts. **f**, Features and conditions used to define CTCF loops. **g**, Quantification of CTCF loops with Cohesin (Coh.), depending on the different configurations in each allele. **h**, Contact map illustrating an allele-specific CTCF loop on Chr17. The track for CTCF orientation shows the directionality of CTCF motifs. Purple arrows point to CTCF motifs with convergent orientation involved in the CTCF loop with CAST specific Rad21 peak in one of the anchors.

We first asked whether the selected E-ASE gene contacts are preferentially established from the most– or least-expressing allele. Similar numbers of E-ASE gene contacts were found to occur in the most or least expressed allele, but rarely in both alleles, suggesting that allele-specific E-ASE contacts can alternatively coincide with the expression of the active allele or with its repression (**Figure 5b**; Suppl. Fig. 7b). Regardless of whether the strong E-ASE contact occurs in the haplotype where the ASE is most or least expressed, we found increased decompaction of the most-expressed allele involved in a strong E-ASE gene contact, by comparing the WDF of the 150 kb genomic regions centered on the ASE gene promoters (**Figure 5c**). These results show that enhancers can contact their putative target genes independently of their compaction or expression state, and confirm that allele-specific expression coincides with increased local decompaction of the expressed genomic region. For example, strong E-ASE gene contacts that coincide with expression of the active allele are in line with models of increased gene expression driven by increased E-P contacts (Carter *et al*, 2002; Simonis *et al*, 2006; Noordermeer *et al*, 2011; Bartman *et al*, 2016). For example, the genes *Fuom* and *Spef1l*, are two CAST ASE genes which establish strong CAST contacts between themselves and enhancer-containing windows spanning a >1 Mb genomic region which is contained within the same compartment A (**Figure 5d**). In contrast, the S129 haplotype is characterized by fewer strong contacts across the whole region, and the presence of a S129-specific compartment B and a LAD interspersing the two CAST ASE genes. WDF measurements show the higher decompaction of the whole region in CAST than S129 haplotypes. We also found examples of loss of strong E-ASE gene contacts in the most active allele, in line with enhancer mechanisms where increased transcriptional activity coincides with loss of E-P contacts (Benabdallah *et al*, 2019). For example, gene *Zfp146* is a CAST ASE gene which establishes an E-ASE strong contact in the S129 allele, spanning 1.7 Mb. Other CAST genes that form strong E-ASE contacts in S129 are *Sirt2* and *Zfp74*, which contact each other (**Figure 5e**). As previously, we find lower WDF in the silent S129 haplotype than CAST haplotype indicating that ASE expression is associated with increased decompaction in the expressing allele.

These results show that allele-specific expression can coincide alternatively with strong allele-specific E-ASE contacts or with loss of strong E-ASE contacts. Irrespective of whether the proximity to putative regulatory regions occurs in the active or repressed state, the most expressed allele is characterized by increased local chromatin decondensation, which may relate to the formation of transcriptional condensates (Cramer, 2019)

### ASE gene contacts include CTCF loops

Finally, we searched for strong ASE contacts anchored by CTCF and RAD21 occupancy, which contained CTCF motifs in convergent orientation and were less than 2 Mb apart (**Figure 5f**; Suppl. Fig. 7c). We found a small number of CTCF loops involving ASE genes (**Figure 5g**), for example, for *Camk1d*, *Gnas* and *H2-Q2* genes (the table of differential contacts and features is available in GSE254717). *H2-Q2* is a CAST ASE gene within the Major Histocompatibility region which is involved in a strong CAST-specific CTCF loop with S129 ASE gene H2-K1 (**Figure 5h**). In total, four histocompatibility genes are ASE genes: *C2* and *H2-Q2* are CAST and *H2-K1* and *H2-D1* are S129 ASE genes. The contact is mediated by two common CTCF peaks with convergent orientation and two cohesin peaks, in which the peak in the right anchor is CAST specific (purple arrows). The CTCF-mediated loop in the CAST allele may favor the expression of *H2-Q2* but not of *H2-K1* in the CAST allele, while the absence of CTCF loop in the S129 correlates with S129 expression of *H2-K1* but not *H2-Q2*. These results suggest that some CTCF loops may be involved in ASE imbalance. However, we observe that these genomic regions are also under Polycomb regulation, for example in specialized cells such as in oligodendroglia (Meijer *et al*, 2022). Histocompatibility genes are also marked by Polycomb histone marks in mESCs and throughout different stages of differentiation of mESCs to neuronal lineages (Ferrai *et al*, 2017). Taken together, these examples showcase the complex interplay between different mechanisms of chromatin and gene regulation and the challenges in the interpretation of the extensive allele-specific differences in 3D genome structure.

## Discussion

Many genes are expressed with allelic imbalance due to a combination of genetic differences between the two chromosome copies, and parental-specific epigenetic mechanisms often attributed to Polycomb repression or DNA methylation (Ohishi *et al*, 2019; Savol *et al*, 2017; Lappalainen *et al*, 2013; Crowley *et al*, 2015; Marion-Poll *et al*, 2021; Inoue *et al*, 2017; Santini *et al*, 2021). Hi-C-based mapping of chromatin contacts showed extensive folding differences between the active and inactive chromosome X copies, which have led to a deeper understanding of the underlying mechanisms of gene repression and escape (Giorgetti *et al*, 2016; Tan *et al*, 2018). In contrast, few differences in 3D chromatin structure have been reported in autosomes based on ligation-dependent methods, including Hi-C, 4C, Dip-C (Llères *et al*, 2019; Rao *et al*, 2014; Han *et al*, 2020). However, ligation-dependent methods combined with standard read length sequencing can only assign 25% of all sequenced ligation events to a specific haplotype, due to the sparsity of SNPs in the genome and the requirement for SNP presence on both sides of the ligation event (Rivera-Mulia *et al*, 2018). In specific mouse crosses where high SNP densities have been achieved (e.g. 1/75 nucleotides; Giorgetti *et al*, 2016), the maximum fraction of phased ligation events are capped at one third of all sequenced ligation events. In human cells, with heterozygous SNP densities closer to 1/2,000 nucleotides, the phasing problem becomes even more demanding and leads to reports of even fewer 3D genome structural differences (Rao *et al*, 2014). These difficulties have been discussed and currently motivate the development of imputation or machine learning approaches that extrapolate unphased events (Miller & Adjeroh, 2024).

In GAM technology, chromatin contacts are inferred by spatial sampling chromosome structure through slicing nuclei in thin slices, and sequencing of the genomic content of each slice (Beagrie *et al*, 2017). Chromatin contacts are measured from the co-segregation of genomic windows in single thin nuclear slices. As the length of each window is typically 10-50 kb, each window typically contains many nucleotide polymorphisms. For example, in hybrid F123 mESCs with average SNP density of 1/124 nucleotides on average, the genotype-specificity of the average 50 kb window can be robustly detected from the sequencing of any combination of an average of 385 SNPs. The GAM sampling process therefore makes the phasing of genomic windows in highly efficient, yielding about 75% of all detected genomic windows in F123 mESC GAM datasets.

In this study, we collected the largest GAM dataset to date from a hybrid mESC line, F123, which has been extensively studied and for which many orthogonal datasets were publicly available (Peric-Hupkes *et al*, 2010; Juric *et al*, 2019; Kubo *et al*, 2021; Li *et al*, 2019). We developed novel pipelines to accurately align reads simultaneously to two different haplotypes, and GAM-phaser to phase GAM data. Phased GAM data revealed an unprecedented level of structural differences between autosomes, at all scales of 3D genome structure and across all autosomes, demonstrating the power of window-based approaches to map haplotype specific differences in chromatin structure.

To begin interpreting the origins and roles of allele-specific chromatin structure, we mapped ASE imbalance using total RNA-seq in the F123 mESC line, and found approximately 2,200 ASE genes, of which 193 are monoallelically expressed. Many ASE genes are housekeeping, with roles in metabolism and signaling, and enriched for genes encoding for ribosomal subunits, as previously reported in other systems, but also histone genes. By mapping the occupancy of Pol2-S5p, Pol2-S2p and H3K27me3 in F123 mESCs, we found that ASE genes often have features of bivalent chromatin and mixed Polycomb-Active promoters states, previously characterized in different mESC lines and throughout neuronal differentiation (Brookes *et al*, 2012; Ferrai *et al*, 2017). A role for Polycomb and mixed allele-specific bivalent promoter states in allele-specific gene expression was also previously reported in mouse embryonic fibroblasts (Savol *et al*, 2017). Amongst the ASE genes with Polycomb-Active promoter states, we found 25 ASE histone genes, 19 of them located within the Hist1 cluster.

As previously reported in other tissues and species (Garcia *et al*, 2014; Crowley *et al*, 2015), ASE genes tend to be clustered in the genome. Some genomic regions have mostly paternal expressed genes, or mostly maternal, but many regions also have both, suggesting that different combinations of genes are expressed from the two chromosome copies. Regardless of their relative genomic positioning, ASE genes are always found in gene-dense regions, intermingled with, or close to, biallelic genes, suggesting that the repression of ASE genes in one allele is likely specific of gene and genomic neighborhood, and not trivially related with 3D chromatin structure. The presence of ASE genes embedded in genomic neighborhoods containing biallelic expressed genes made the exploration of allele-specific 3D chromatin structure especially challenging, namely in understanding why ASE genes are sensitive to local repression/activation mechanisms or contribute to the allele-specific chromatin structure, while the neighboring biallelic genes are seemingly refractory.

The extensive differences in 3D genome structure between the two copies of each chromosome were observed at all levels of 3D genome organization, both locally and spanning large genomic regions. Haplotype-specific GAM data detected differences in compartment A/B annotation across 18% of the genome, a much larger proportion than previously reported using Hi-C in cells with similar SNP density (4% in mouse cell data; Han *et al*, 2020). We also found extensive differences in chromatin insulation at the level of topologically associating domains (TADs). Of all TAD borders detected in both CAST and S129 alleles, the majority (59%) are allele-specific. Haplotype-specific TAD borders are nevertheless characterized by an enrichment in CTCF and cohesin occupancy, albeit to a lower extent than common TAD borders present in both haplotypes. Interestingly, we found that allele-specific CTCF occupancy on chromatin was generally rare (6.5% of all CTCF peaks were allele specific compared with 78% biallelic CTFC peaks). Interestingly, haplotype-specific CTCF peaks occurred most often inside TADs, suggesting that haplotype-specific TAD border formation is not a trivial consequence of mechanisms based on haplotype-specific CTCF occupancy. We also found that TAD organization was related to ASE gene clustering. Remarkably, ASE genes are present in only half of all TADs, and their presence coincides with an increased presence of H3K27me3 occupancy. By assessing chromatin compaction directly from GAM data, based on the detection frequency of each genomic region, we found that Polycomb occupancy coincides genome-wide with increased compaction of the least expressed allele, adding to previous *in vitro* and *in vivo* observations (Nichols *et al*, 2020; Barbieri *et al*, 2017). In contrast, allele-specific methylation occurs at a minority of genes, as suggested previously (Kerkel *et al*, 2008), especially at monoallelic genes, and is not a major feature of ASE imbalance. However, when searching for putative transcription factor binding sites at ASE promoters, genic and intergenic regions, we discovered that a single transcription factor, ZFP57, is specifically enriched at the promoters of CAST ASE genes, suggesting a role for haplotype or parental-specific effects in the occupancy and regulation of ASEs in mESCs.

Amongst the clustered ASEs associated with Polycomb chromatin occupancy, we explored in more detail the Hist1 locus, which contains 19 ASE histone genes in F123 mESCs. We found that the Hist1 cluster contacts are highly haplotype specific and especially predominant in the S129 allele where fewer ASE histone genes are expressed. We also noticed that the Hist1 cluster is abundantly covered by H3K27me3-marked chromatin (25% of 50 kb windows are positive for H3K27me3 peaks), and that many of the haplotype-specific contacts in the S129 genome occur between genomic windows marked by Polycomb occupancy. To functionally test a role for Polycomb repression in histone gene downregulation, we took advantage of available conditional knockouts of the two major enzymatic subunits of Polycomb Repressor Complexes, PRC1 (*Ring1b*) and PRC2 (*Ezh2*). Tamoxifen-induced knockouts of *Ezh2* or *Ring1b* led to upregulation of histone protein levels, showing that histone genes are direct targets of Polycomb repression mechanisms. Further work will be necessary to investigate at the single cell level how the increased clustering of the Hist1 locus in the S129 genome relates to the lower expression of specific histone genes in each haplotype, and how allele-specific repression of specific histone genes relates to the cell cycle, the histone locus body or Polycomb bodies (Ghule *et al*, 2008; Nizami *et al*, 2010; Quinodoz *et al*, 2018).

Even though we explored other chromatin features, most were typically biallelic. Nevertheless, deeper investigation of allele-specific enhancer contacts highlighted potential candidate regulatory regions strongly contacting specific ASE genes in one specific allele. Allele-specific CTCF-loops were also rare but occasionally associated with ASE genes. Amongst the genes involved in haplotype-specific CTCF loops, we found immune system genes. Immune system genes have been reported to be ASE in other biological systems, including in F1 crosses between goats and Ibex, or between humans and Neanderthals, and associated with disease (Yang *et al*, 2022, McCoy *et al*, 2017). Moreover, histocompatibility genes are susceptible to cis regulation variants (Gutierrez-Arcelus *et al*, 2020). The observation that histocompatibility genes form highly haplotype-specific contacts in a haplotype– or parental-specific manner indicates a role for 3D genome structure in the diversity of major histocompatibility complexes and the capacity of the immune system evolution, which requires further work in relevant biological systems (Sommer, 2005).

Overall, the observation of a variety of chromatin regulatory mechanisms connected with ASE imbalance highlights that ASE imbalance is tuned by combinations of different mechanisms and that regulation is highly gene specific (Crowley *et al*, 2015; Marion-Poll *et al*, 2021). These findings also demonstrate the value of haplotype-specific 3D genome structure to help address mechanisms of disease due to genetic variation or epigenetic deregulation of genes. Future questions and limitations of the present study are the contribution of parental versus genetic sequence effects, which can be addressed by mapping allele-specific differences in the alternative cross (S129xCAST) and using other genotypes. Further efforts are required to understand the stability and evolution of allele-specific chromatin structures in differentiation and in different cell lineages. The detected differences between CAST/paternal and S129/maternal phasing of local features open new questions about parental-specific epigenetic mechanisms acting on allelic expression imbalance, which require in-depth further study. Further work is also necessary to enable the phasing of GAM from human samples, which are characterized by ten times lower SNP densities than F123 mESCs. Finally, it is still an open question to what extent phasing the allele-specific topology of the two chromosome copies can help to interpret genome sequence and predict its effects on gene (de)regulation, towards a deeper understanding of genome biology and gene regulation mechanisms.

## Material and Methods

### Cell culture

F123 mESCs (a male, hybrid cell line, derived from a F1 S129/Jae and Cast mouse cross) (Gribnau et al., 2003). Cells were cultured in a layer of mitotically inactivated feeder murine embryonic fibroblasts under standard conditions (DMEM, supplemented with 15 % KSR, 1x Glutamax, 10 mM non-essential amino acids, 50 µM beta-mercaptoethanol, 1,000 U/ml LIF). Before harvesting, mESCs were passaged onto feeder-free 0.1% gelatin coated plates for at least 2 passages to remove feeder cells. As feeder removal results in reduced levels of LIF in the culture, the LIF concentration in the media was doubled when the cells were in feeder-free culture conditions. Cells were harvested after approximately 48 h at 70 – 80 % confluency.

ESC-ERT2 Ring1A−/− cells (Stock *et al*, 2007) were maintained in an undifferentiated state by co-culture on mitomycin-inactivated mouse embryonic fibroblasts on 0.1% gelatin-coated flasks in DMEM supplemented with non-essential amino acids, 2 mM L-glutamine, 0.1 mM 2-mercaptoethanol (all from Gibco), 20% FCS (Autogen Bioclear, Calne, UK) and 1,000 U/ml of leukemia inhibitory factor (ESGRO-LIF, Chemicon/Millipore). For the Ring1B conditional deletion, ESC-ERT2 cells were plated feeder-free on gelatin-coated plates 12 h before supplementing the medium with 800 nM 4-hydroxytamoxifen (H7904, Sigma, Poole, UK), and grown for 48h.

ESC-Ezh2-1.3 cells (Pereira *et al*, 2010) were maintained in an undifferentiated state by co-culture on mitomycin-inactivated mouse embryonic fibroblasts on 0.1% gelatin-coated flasks in Knockout DMEM supplemented with non-essential amino acids, 2 mM L-glutamine, 0.1 mM 2-mercaptoethanol (all from Gibco), 20% FCS (Autogen Bioclear, Calne, UK), 5% Knockout Serum Replacement (Invitrogen), and 1,000 U/ml of leukemia inhibitory factor (ESGRO-LIF, Chemicon/Millipore). For the Ezh2 conditional deletion, ESC-Ezh2-1.3 cells were plated feeder-free on gelatin-coated plates 12 h before supplementing the medium with 800 nM 4-hydroxytamoxifen (H7904, Sigma, Poole, UK), and grown for 96h, including replating at 48h.

### Total RNA sequencing

Total RNA was extracted from F123 mESCs using TRIzol Reagent (Invitrogen, Cat# 15596026) following manufacturer’s instructions. Total RNA was analyzed on the Bioanalyzer using the Agilent RNA 6000 Nano Kit to ensure intact, non-degraded RNA presence and was subsequently treated with TURBO DNase I (Ambion, Cat# AM1907). Total RNA-seq libraries were generated from 1 μg of DNase-treated RNA using the TruSeq Stranded total RNA library preparation kit (Illumina, Cat# 15031048) according to the manufacturer’s instructions. Samples were pooled and paired-end (75 bp) sequenced using an Illumina NextSeq500/550 sequencer, following the manufacturer’s instructions.

### Genome Architecture Mapping (GAM)

Fixation of F123 mESCs was performed as described previously (Branco and Pombo, 2006). Briefly, mESCs were grown to 70 % confluency, media was removed, and cells were fixed in 4% and 8% paraformaldehyde in 250 mM HEPES-NaOH (pH 7.6; 10 min and 2 h, respectively), gently scrapped, and softly pelleted, before embedding (>2h) in saturated 2.1 M sucrose in PBS and frozen in liquid nitrogen on copper sample holders. Frozen mESCs samples were stored indefinitely in liquid nitrogen.

Ultrathin cryosections were cut with a glass knife using an ultracryomicrotome (Leica Biosystems, EM UC7) at ∼230 nm thickness, and transferred to UV-irradiated PEN membrane steel frame slides 4.0 µm (Leica Microsystems, 11600289) for laser microdissection. Before laser microdissection, cryosections were washed in sterile-filtered 1x PBS (3 times, 5 min each) to remove the sucrose, sterile-filtered water (3 times, 5 min each), and stained with sterile-filtered 1 % (w/v) cresyl violet (Sigma-Aldrich, C5042) in water for 10 min, followed by 2 washes with water (30 s each). Individual nuclear profiles (NPs) were isolated by laser microdissection using a laser microdissection microscope (Leica Microsystems, LMD7000). Cut NPs were collected in a PCR Cap Strip filled with opaque adhesive material (Carl Zeiss Microscopy, 415190-9161-000). For each collection day, 1 or 2 caps were left empty and taken through the whole genome amplification (WGA) and sequencing process as a negative control for quality control purposes (labeled as “0NP” samples in SI Table 10).

Whole Genome Amplification (WGA) was performed as described previously (Winick-Ng *et al*, 2021) with minor modifications. Briefly, DNA was extracted from NPs at 60^0^C in the lysis buffer (20 mM Tris-HCl pH 8.0, 1.4 mM EDTA, 560 mM guanidinium-HCl, 3.5% Tween-20, 0.35% Triton X-100) containing 0.75 units/ml Qiagen protease (Qiagen, 19155). After 24h of DNA extraction, the protease was heat inactivated at 75 °C for 30 min and the extracted DNA was amplified via two rounds of PCR. At-first quasi-linear amplification was performed with random hexamer GAT-7N primers with an adaptor sequence. The lysis buffer containing the extracted genomic DNA was mixed with 2x DeepVent mix buffer (2x Thermo polymerase buffer (10x), 400 µm dNTPs, 4 mM MgSO_4_ in ultrapure DNase free water), 0.5 µM GAT-7N primers (5′-GTG AGT GAT GGT TGA GGT AGT GTG GAG NNN NNN N) and 2 units/µl DeepVent^®^ (exo-) DNA polymerase (New England Biolabs, M0259L) and incubated for 11 cycles in the BioRad thermocycler. The second exponential PCR amplification was performed in presence of 1x DeepVent mix, 10 mM dNTPs, 0.4 µM GAM-COM primers (5′-GTG AGT GAT GGT TGA GGT AGT GTG GAG) and 2 units/µl DeepVent (exo-) DNA polymerase in the programmable thermal cycler for 26 cycles.

### GAM library preparation and high-throughput sequencing

After whole genome amplification, the DNA WGA product was purified with SPRI beads (1.7x) The DNA concentration of each sample was quantified using the Quant-iT® PicoGreen dsDNA assay kit (Invitrogen #P7589). Genomic sequencing library was prepared from 1 ng of purified DNA using the Illumina Nextera XT library preparation kit (Illumina #FC-131-1096), following the manufacturer’s instructions or with a reduced volume of reagents to 20%. The library preparation step was done either manually or using TTP Mosquito HV liquid handling system, as specified in SI Table 10 After the library preparation, DNA was again purified with in-house SPRI beads (1.7x) and equal amounts of DNA from each sample was pooled together (up to 196 samples) for the sequencing. The final pool of libraries was purified two more times with SPRI beads (1.7x) and analyzed using DNA High Sensitivity on-chip electrophoresis on an Agilent 2100 Bioanalyzer. The samples were sequenced on an Illumina NextSeq500/550 sequencer as single-end 75 bp reads, according to manufacturer’s instructions.

### SNP calling in F123 hybrid (N-masked genome generation)

The high SNP density of the F123 genome was used to phase the reads from sequenced GAM libraries to the maternal and paternal haplotypes. For generating haplotype specific calls for the hybrid F123 (CAST×S129) cells, the parental genome sequencing data from publicly available databases was used. The genome sequence of *Mus musculus castaneus* was downloaded from the European Nucleotide Archive (accession number ERP000042). *Mus musculus musculus S129/SvJae* genome sequence data was downloaded from the Sequence Read Archive (accession number SRX037820). Read trimming was performed using Cutadapt (https://cutadapt.readthedocs.io/en/stable/, Martin, 2011) and mapped the reads to the mm10 genome assembly using the Burrows-Wheeler Aligner (https://bio-bwa.sourceforge.net, Li & Durbin, 2010). SNP location and sequence were identified using bcftools (http://samtools.github.io/bcftools/bcftools.html). SNPs that had less than 5 reads per SNP and quality below 30 were excluded from the analysis.

### Phasing of GAM data with theGAM-Phaser pipeline

GAM-Phaser is a pipeline developed here for GAM data phasing, summarized in Suppl. Fig 1b. GAM-Phaser takes as input a VCF file (file containing position of SNPs) and raw GAM sequencing data (fastq files) and outputs haplotype-specific GAM window segregation tables for each haplotype considered. GAM-phaser takes advantage of the existing SNPsplit package (Krueger & Andrews, 2016) to mask high-quality paternal and maternal SNPs with N-character in a genome. The mm10 reference genome assembly was used (Dec. 2011, GRCm38/mm10).

GAM-phaser generates an N-masked genome using the information about SNPs genomic coordinates and the reference genome assembly via SNPsplit package. At the next step, raw GAM sequencing data are mapped to the N-masked genome using default parameters of bowtie2 (version 2.3.4.3) (Langmead & Salzberg, 2012). The reads mapped to the N-masked genome are checked for the presence or absence of a SNP, and sorted to the haplotype specific bam-files with SNPsplit package. Next, the genome is split into equal-sized windows and the coverage of all reads, CAST-phased reads and S129-phased reads is computed using bedtools for all collected F123 GAM libraries (Quinlan & Hall, 2010). Afterwards, the optimal threshold between the sequencing noise and the signal is determined separately for each GAM sample of the total dataset. The optimal threshold of nucleotide coverage for calling positive windows is calculated as the lowest coverage per bin that gives the highest percent of windows that have at least one neighboring positive window on at least one side). Windows are phased to the CAST haplotype when the number of nucleotides covered by the reads containing CAST SNPs is higher or equal to the optimal threshold, and to the S129 allele when the number of nucleotides covered by the reads containing maternal SNPs is higher or equal to the optimal threshold.

### Quality control of GAM samples

After read mapping and positive window calling, the quality of each GAM sample in the dataset collected was accessed to ensure that the laser microdissection, DNA extraction and subsequent experimental steps, were successful. Quality control metrics calculated for each GAM sample include the number of uniquely mapped reads to the mouse genome, the percentage of orphan windows (windows without at least one neighbor) and the percent of total genome coverage. To exclude potentially cross-contaminated samples, Jaccard similarity index was calculated between the sequences of positive and negative windows from all GAM samples that were processed together on the same 96-well plate, as previously reported (Winick-Ng 2017). Samples with a Jaccard similarity index > 0.4 were excluded from the data analysis as potentially contaminated. A sample was considered to be a good quality if it had < 60% orphan windows, > 50,000 uniquely mapped reads and did not appear as cross-contaminated (Suppl. Fig. 8a). The detailed quality metrics for all samples are provided in SI Table 10.

Most (90.51%) of the collected NPs passed quality control. The final GAM dataset was composed of 3,707 high quality nuclear profiles (NPs), and sampled from two biological replicates: 863 NPs were collected in 3NP mode (549 from replicate 1 and 314 from replicate 2), while 1,122 NPs were collected in 1NP mode (from replicate 1) and combined to 3NP *in silico (see* SI Table 11 and SI Table 12), as described previously (Winick-Ng *et al*, 2021, Beagrie *et al*, 2023) (Suppl. Fig. 1a).

### Determining resolution of pairwise co-segregation matrices

To assess the quality of genome sampling, the distribution of raw co-segregation events for all intra-chromosomal pairs of genomic windows was compared to the standard Poisson distribution, at different resolution(s) and genomic distance(s) using a non-parametric Kendall rank correlation coefficient. The calculation of raw cosegregation events was followed by Yeo–Johnson power transformation. Standard Poisson distribution was computed using the mean and the standard deviation derived from the distribution of the real co-segregation events at each tested resolution(s) and genomic distance(s). Kendall’s τ correlation coefficient ≥ 0.95 was considered as the indication of good quality of genome sampling at specified resolution and genomic distance.

### GAM data normalization

Raw cosegregation GAM matrices were normalized using normalized pointwise mutual information (NPMI) for all pairs of windows genome-wide, as previously described (Winick-Ng *et al*., 2021). NPMI describes the difference between the probability of a pair of genomic windows being found in the same NP given both their joint distribution and their individual distributions across all NPs. For visualization purposes, scale bars were adjusted to a range 0 and the NPMI value corresponding to the 99th percentile of all NPMI values for each genomic region displayed.

### Window Detection Frequency calculation

Window detection efficiency (WDF) is a GAM specific parameter representing the amount of times a specific window is captured in the whole GAM dataset on average (Beagrie *et al*, 2017). It is calculated from the GAM segregation tables, by dividing the total number of times that each genomic window is present in the whole dataset by the total number of samples in the dataset:

WDF = SUM(Positive Window x) / SUM(samples)

WDF was calculated from the combined 3DN-GAM segregation tables at 50 kb resolution, separately for CAST and S129 alleles (SI Table 8).

### Identification of undersampled regions in GAM contact matrices

The WDF of genomic windows was also used to exclude genomic windows which are insufficiently sampled in the GAM process from further analyses. WDF data follows a normal distribution in the genome. To detect outliers, a smoothing algorithm was applied to the WDF values per chromosome in stretches of eleven 50 kb genomic windows. Next, normalized delta (ND) was calculate for each window, according to

ND = (raw_Signal – smoothed_Signal)/smoothed_Signal

If the ND is larger than a fold change of 5, the window is removed from the final dataset.

Next, the four adjacent windows (2 upstream and 2 downstream) to the window being removed were also removed, to ensure good quality of sampling in the final GAM data used for further analyses. Finally, genomic bins with an average mappability score below 0.2 are removed. Genome mappability for mm10 mouse genome assembly was computed using GEM-Tools suite (Marco-Sola *et al*, 2012) setting read length to 75 nucleotides. The mean mappability score was computed for each genomic bin with bigWigAverageOverBed utility from Encode.

### Non-redundant gene list selection

The most expressed isoform for each gene was identified using the same strategy as in Ferrai *et al* with some modifications (2017). Briefly, a complete expression analysis table containing 39,261 unique genes and 88,437 isoforms was considered. Almost 20,000 genes (n=19,003) had a single annotated isoform. For the remaining 20,258 genes, a single isoform was selected based on the following criteria: (i) gene isoform with the highest amount of reads for Pol2-S5p in 2 kb window centered on the TSS (14,051 genes); (ii) if ambiguity was still present, the gene isoform with the highest amount of reads for Pol2-S7p in the 2 kb window centered on the TSS was selected (1,623 genes); (iii) if ambiguity was still present, the longest gene isoform was selected (3,827 genes); (iv) if ambiguity was still present, a random annotated isoform was selected (757 genes).

### Promoter state classification

To classify gene promoter states, we followed the same strategy as in (Ferrai *et al*, 2017) with some modifications. Briefly, gene promoters were considered positive for Pol2-S5p, Pol2-S7p, or H3K27me3 when: (i) the 2 kb windows centered on the TSS overlapped with a region enriched for the mark, and (ii) the amount of reads in the TSS window was above a threshold. The threshold was defined as the 5th percentile of the distribution of reads in the TSS window of positive genes. Overlapping genes (3,558) and genes whose TSSs were in close proximity (6,855) were excluded from the classification. In total, we identified 6,435 active genes, 12,968 inactive genes, 1,716 PRC repressed and 5,082 Polycomb-Active genes.

### RNA-seq data analysis

RNAseq data from F123 was processed for standard and allele-specific gene expression analysis. The quality of the paired-end RNA sequencing reads was verified using FASTQC (http://www.bioinformatics.babraham.ac.uk/projects/fastqc). No reads needed to be trimmed or removed due to quality concerns. The paired-end reads derived from RNA sequencing were mapped to the most recent mouse reference genome assembly mm10 (GRCm38.p6) using STAR (version 2.7.2c) (Dobin *et al*, 2012) under consideration of the current mm10 annotation (downloaded from ensemble: ftp://ftp.ensembl.org/pub/current_gtf/mus_musculus/Mus_musculus.GRCm38.98.gtf.gz) and available information of genomic variants in the mm10 F123 genome (described in SNP calling in F123 hybrid). Following recommendations about best practices for data processing in allelic expression analysis (Castel *et al.,* 2015), duplicate reads were removed from the data using Picard MarkDuplicates (version 2.21.1: https://software.broadinstitute.org/gatk/documentation/tooldocs/4.1.3.0/picard_sam_markdupl <u>icates_MarkDuplicates.php</u>). Default options were used with the exception of REMOVE_DUPLICATES = TRUE. To quantify the overall expression of genes, mapped reads overlapping exons and introns were assigned to the respective genes and summarized as gene specific count values using HTSeq-count (Anders *et al*, 2014). The use of HTSeq-counts to generate gene level read count values is recommended by the gold standard tool used for differential gene expression analysis DESeq2 (Love *et al*, 2014). Options were set to count reads overlapping exons and introns of genes, accounting for the paired end nature of reads, only considering primary alignments and the default minimal alignment quality of 10. The same annotation file was used as described before in the read mapping step.

Subsequently TPM values were calculated by normalizing count values for gene length and library size. To differentiate between the expression of genes located on the two parental alleles, reads that overlap heterozygous genomic variants were counted in an allele-specific manner. Reads overlapping those heterozygous variants located within exons and introns of genes were counted using GATK ASEReadCounter (The Genome Analysis Toolkit (GATK) version 4.1.3.0: https://software.broadinstitute.org/gatk/documentation/tooldocs/4.1.3.0/org_broadinstitute_he <u>llbender_tools_walkers_rnaseq_ASEReadCounter.php</u>). Subsequently, only genomic variants within regions of high mappability and with a minimum total coverage of 20 reads were considered to reduce the risk of introduced biases. In case multiple genomic variants were present within the same gene, the counts were aggregated over the gene in an allele specific manner using the available haplotype information described above in the read mapping step. Aggregated counts were tested for significant allele specific expression differences (binomial test vs 0.5) and the false discovery rate was controlled for by correcting resulting p-values for multiple testing using the Benjamini and Hochberg method. Genes were defined as differentially expressed by an adjusted p-value ≤0.05, and a fold change (log2) ≥ 1 between reads mapping to CAST and S129. The ASE ratio was calculated as the ratio of read counts supporting the CAST haplotype and total read count. The Log2 fold change was defined as the log-scaled ratio of reads supporting the CAST haplotype divided by the read count observed in the S129 haplotype.

### Gene Ontology (GO) enrichment

GO enrichment analysis of genes with allele-specific expression was performed using Web Gestalt (https://www.webgestalt.org/). All expressed genes were used as the background universe. Over representation analysis was performed selecting Gene Ontology as a Functional database in the website.

### Insulation scores calculation and topological domain boundary calling

TAD calling was performed by calculating insulation scores in NPMI GAM contact matrices at 50 kb resolution, as previously described (Winick-Ng *et al*, 2021), using the insulation square method. The insulation score was computed with insulation square sizes ranging from 100 to 1,000□kb for the unphased matrices and each haplotype. TAD borders were called using a 400 kb insulation square size and based on local minima of the insulation score with one genomic bin added on each side.

### Allele quantification with cryo-FISH

We obtained the source cryo-FISH data for the detection of 40 kb genomic regions containing the Hoxb1 or Hoxb13 genes performed in mESCs clone 46C, which reports for each nuclear slice, whether 1 or 2 copies of each locus are present (Barbieri *et al*, 2017). We counted the number of sections that contained both alleles and divided for the number of sections that contained one or two alleles. We performed this analysis for two channels: the green that corresponded to Hoxb1 locus and the red that corresponded to the Hoxb13 locus.

### Identification of compartments A and B

Compartments were calculated using 100 kb resolution co-segregation matrices. In brief, each chromosome was represented as a matrix of observed interactions O(i,j) between locus i and locus j. We then calculated the expected interactions E(i,j) matrix, where each pair of genomic windows is the mean number of contacts with the same distance between i and j. A matrix of observed over expected values O/E(i,j) was produced by dividing O by E. A correlation matrix C(i,j) was calculated between column i and column j of the O/E matrix. PCA was performed for the first three components on matrix C before extracting the component with the best correlation to GC content. Loci with PCA eigenvector values with the same sign that correlate best with GC content were called A compartments, whereas regions with the opposite sign were B compartments. Eigenvector values on the same chromosome in compartment□A were normalized from 0 to 1, whereas values on the same chromosome in compartment□B were normalized from −1 to 0.

### Identification of allele-specific contacts

Allele-specific contacts were identified using a previously developed pipeline for finding differential contacts between two contact maps (Beagrie et al, 2023; Winick-Ng et al, 2021) with some adjustments to adapt for the allelic setting. Following the removal of undersampled regions and setting a maximum contact distance of 50 Mb, each chromosomal contact matrix at 50 kb resolution from CAST and S129 NPMI was transformed into their z-scores equivalent, by adjusting for the mean and variance across all contact distances. Next, the difference between both alleles was computed by subtracting normalized S129 contacts from CAST contacts (delta-z-score=CAST-S129). Finally, contacts with delta z-score below –1 or above 1 and NPMI intensities above 0.3 in either of the two maps were selected as S129-specific or CAST-specific contacts, respectively, to focus the subsequent analyses on the strongest contacts.

### Identification of strong allelic contacts

Strong allelic contacts represent the highest values on each chromosome of each allele. In contrast to allele-specific contacts which are specific to one haplotype, strong contacts are not informed by the alternative allele and, in consequence, strong CAST contacts can also be strong in the S129 allele. The strongest contacts in the CAST allele and S129 allele were extracted using an NPMI score >0.3 and a z-score >2.0 in the distance-normalized matrices from each haplotype, respectively (see Identification of differential contacts).

### Distance decay and derivatives calculation

Decay plots and momentum curves (Abdennur *et al*, 2022) or S129 and CAST contacts maps were calculated using the mean contact intensity over distance displayed at logarithmic scale (log10). Momentum curves were obtained from the *ksmooth* R function using a Normal kernel with bandwidth of 0.3. The slope values in CAST or S129 contact decay are based on derivatives obtained from the difference between observed mean intensity scores at equidistant breakpoints, set at log10-scaled distance intervals of 0.1.

### ATAC-seq data mapping, processing, QC and phasing

ATAC-seq reads were mapped, quality controlled, and split into their respective genomes using SNPsplit. Then, peaks were called with ChromA (https://github.com/marianogabitto/ChromA, Gabitto *et al*, 2020). D score was calculated for each peak, as a measure of their allelic imbalance in order to assign allele-specific peaks followed by a permutation test to assess their significance (Xu *et al*, 2017). Finally, a stringent filtering was applied to identify allele-specific peaks, requiring both biological replicates to have a Dscore between –0.3 and –0.3 (reads ratio score), a minimum of ten reads in the peak, and a p-value <0.01 in the permutation test, after FDR correction according to Benjamini and Hochberg.

### Motif calling in ATAC-seq peaks

First, *annotatePeaks.pl* script from the Homer tools suite is run in the CAST specific, S129 specific, common or unphased ATAC peaks, to classify them depending on their genomic position. Then for each type, the closest gene was identified, which is the most likely to be the target gene. Finally, for each of these groups, *findMotifsGenome.pl* is run to find the enriched motifs. Q-value of ≤ 0.05 and p-value of ≤ 0.001 was used as cutoff for enriched motifs.

### ChIP-seq data collection, QC, mapping and processing

Chromatin immunoprecipitation experiments were performed as previously described (Brookes *et al*, 2012; Ferrai *et al*, 2017). Pol2-S5p was detected with mouse antibodies CTD4H8 clone (BioLegend, Cat# 904001); Pol2-S7p with rat antibodies 4E12 clone (Chapman *et al*, 2007; kindly provided by Dirk Eick); Polycomb mark H3K27me3 was detected with rabbit antibodies (Millipore, Cat# 07-449). ChIP-seq libraries were prepared from 10 ng of immunoprecipitated DNA using TruSeq ChIP Library Preparation Kit (Illumina, IP-202-1012) according to the manufacturer’s instructions with minor modifications. Library size was assessed before high-throughput sequencing by Bioanalyzer (Agilent) using the High Sensitivity DNA analysis kit (Agilent, Cat# 5067-4626). ChIP-seq libraries were sequenced 75 bp single-end using Illumina NextSeq500/550 sequencer, according to the manufacturer’s instructions.

### ChIP-seq peak calling and phasing

Raw ChIP-seq reads were mapped to the N-masked genome using default parameters of bowtie2 (version 2.3.4.3) (Langmead & Salzberg, 2012). Genome-wide enriched regions for Pol2-S5p, Pol2-S7p, and H3K27me3 were identified with Bayesian Change-point Model (BCP) peak-finder (Xing *et al*, 2012, default settings). Genome-wide enriched regions for H3K27ac, H3K4me3, CTCF and Rad21 were identified with MACS2 peak finder (Zhang *et al*, 2008; broad peaks, default settings). If two biological replicates were available (CTCF, H3K4me3, H3K27ac, Rad21), the peak calling was performed in each dataset separately and then peaks identified in both datasets were used for further analysis. Next, ChIP-seq reads were phased for all datasets using the SNP-split package and the number of reads in each peak was computed with bedtools coverage (Suppl. Fig. 8b) (Quinlan & Hall, 2010). To classify the peaks as allele specific, the ratio between CAST and S129 allele specific reads was computed for each peak. Peaks that have log2 fold change > 2 were selected as allele specific. Peaks that had <10 SNP-containing reads were excluded from further analysis. Robust identification of allele-specific peaks for broad marks (Pol2-S5p, Pol2-S7p, and H3K27me3) was not possible due to high heterogeneity of the distributions of CAST and S129 specific reads.

### CTCF orientation calling

*annotatePeaks.pl* script was used from the *Homer* suite tools to call the orientation of CTCF motifs in CTCF peaks (http://homer.ucsd.edu/homer/ngs/annotation.html).

### Genome-wide feature co-occurrence

First, the genome is binned in 200 kb bins. For each feature, the number of peaks are counted in each bin, creating a list. Finally, these lists are correlated and a Pearson correlation coefficient is calculated for each comparison.

### Differential DNA methylation

Since bisulfite treatment causes C to T transitions, certain SNPs positions may not be used for allele-specific reads sorting since they might reflect either an allele specific difference or a methylation state. To overcome this limitation, a modified N-masked genome was prepared using Bismark software package (Krueger & Andrews, 2011) and analyzed publicly available whole-genome bisulfite sequencing data (Li *et al*, 2019) with SNPsplit package in WGBS-compatible mode (Krueger & Andrews, 2016). The reads were trimmed using the Trim Galore package using the default settings prior to mapping (Martin, 2011). The methylation calls for every analyzed C were extracted using bismark_methylation_extractor script.

For each allele, CpGs with a methylation percentage higher than 50 were taken for further analysis. Next, the ratio of methylated/unmethylated CpGs in the promoter (±1,000 bp from TSS) of genes bigger than 2,000 bp was calculated. The ratio in the CAST allele was subtracted to the ratio in the S129 allele, giving the differential percentage of possible methylated CpGs. Finally, differentially methylated promoters were those where this differential percentage exceeded the 5th percentile.

### Proteomics

ESC-ERT2 cells grown in DMEM media for SILAC were used as SILAC reference. Cell were cultured in DMEM media lacking L-lysine and L-arginine amino acids, supplemented with 15% knockout serum replacement (KOSR; Invitrogen, #10828), cytokine leukemia inhibitory factor (LIF, Merck, #ESG1107) and heavy amino acid isotopes (L-lysine, +8Da; Cambridge Isotope Laboratories, #CNLM291H; L-arginine +10Da; Cambridge Isotope Laboratories, #CNLM-539H; Bendall *et al*, 2008). Cells were lysed in urea buffer (8M urea, Tris 100 mM, pH 8.25) and sonicated a Bioruptor sonicator (Diagenode), using 10 cycles of sonication (30 s ON, 30 s OFF). After centrifugation to remove debris, protein concentration was measured by Bradford colorimetric assay and 50 µg protein extract were mixed with an equal amount of reference heavy sample. The disulfide bridges of proteins were then reduced in 2 mM DTT for 30 min at 25 °C and successively free cysteines alkylated in 11 mM iodoacetamide for 20 min at room temperature in the dark. LysC digestion was then performed by adding 2 µg of LysC (Wako) to the sample and incubating for 18 h, under gentle shaking at 30 °C. After LysC digestion, the samples were diluted 3 times with 50 mM ammonium bicarbonate, before addition of 7 µl of immobilized trypsin (Applied Biosystems) and incubation for 4 h under rotation at 30 °C. 18 µg of the resulting peptide mixtures were desalted on STAGE Tips (Rappsilber et al, 2002) and the eluates dried and reconstituted to 20 µl of 0.5 % acetic acid in water.

### LC-MS/MS analysis

Five microliters of each sample were injected into the HPLC system (Eksigent) coupled to an Orbitrap Velos mass spectrometer (Thermo). The chromatographic separation using a 240 min gradient ranging from 5% to 45% of solvent B (80% acetonitrile, 0.1% formic acid; solvent A= 5% acetonitrile, 0.1% formic acid). A 30 cm long capillary column (75 µm inner diameter) was packed with 1.8 µm C18 beads (Reprosil-AQ, Dr. Maisch). A tip was generated on one end of the capillary nanospray using a laser puller, allowing fritless packing. The nanospray source was operated with a spray voltage of 1.9 kV and an ion transfer tube temperature of 260 °C. Data were acquired in data dependent mode, with one survey MS scan in the Orbitrap mass analyzer (30,000 resolution at 400 m/z) followed by up to 10 MS/MS scans in the Orbitrap analyzer (15,000 resolution at 400 m/z) on the most intense ions. Once selected for fragmentation, ions were excluded from further selection for 45 s, to increase new sequencing events.

### Proteomics data analysis

Raw data were analyzed using the MaxQuant proteomics pipeline v2.1.3.0 and the built in the Andromeda search engine (Cox & Mann, 2008; Cox *et al*, 2011) with the Uniprot mouse protein database. Carbamidomethylation of cysteines was chosen as fixed modification, oxidation of methionine and acetylation of N-terminus were chosen as variable modifications. Two missed cleavage sites were allowed and peptide tolerance was set to 7 ppm. The search engine peptide assignments were filtered at 1% FDR at both the peptide and protein level. The ‘match between runs’ feature was enabled, ‘second peptide’ feature was enabled, while other parameters were left as default.

## Data availability

Raw fastq sequencing files for all GAM F123 samples, together with non-normalized cosegregation matrices, normalized pair-wised chromatin contact maps, table of contacts with overlapping features and raw GAM segregation tables have been submitted to the GEO repository under accession number GSE254717. Raw and processed data for all GAM F123 samples are also available from the 4DN data portal (https://data.4dnucleome.org/) under accession number 4DNESRQDNG61. Insulation score values of unphased, CAST and S129 alleles are archived in a permanent data repository (Irastorza-Azcarate *et al*., 2024). Raw fastq total transcriptome data from F123 mESCs, together with PM values and ASE gene classifications are available from GEO repository under accession number GSE254675. File with gene expression levels, epigenetic features and classification of gene transcripts is archived in a permanent data repository (Irastorza-Azcarate *et al*., 2024). Raw fastq chromatin occupancy for Pol2-S5p, Pol2-S7p and H3K27me3 in F123 mESCs, together with peak calls for all ChIP-seq datasets produced in this study have been submitted to the GEO repository under accession number GSE254710. Coordinates of phased and unphased peaks for: ATAC, H3K4me3, H3K27ac, CTCF and Rad21 are archived in a permanent data repository (Irastorza-Azcarate *et al*., 2024). List containing transcription factor motif enrichment at CAST or S129 ATAC peaks at promoters, intergenic or genic regions is archived in a permanent data repository (Irastorza-Azcarate *et al*., 2024).

The mass spectrometry proteomics data have been deposited to the ProteomeXchange Consortium via the PRIDE (Perez-Riverol *et al*, 2021) partner repository with the dataset identifier PXD048969.

## Acknowledgements

The authors thank Lonnie Welch, Yingnan Zhang, Luca Fiorillo and Francesco Musella for exploratory data analysis, all laboratory members and the 4D Nucleome consortium for helpful discussions, Dirk Eick for the kind gift of the Pol2-S7p monoclonal antibodies (4E12 clone), and the Genomics Technology Platform and the Protein Production and Characterization Platform, both at the Max Delbrück Center for Molecular Medicine in the Helmholtz Association (MDC), Berlin.

A.P., B.R. and M.N. acknowledge support from the National Institutes of Health Common Fund 4D Nucleome Program grants U54DK107977 and 1UM1HG011585-03.

A.P. and R.F.S. acknowledge support from the Helmholtz Association (Germany) and the DFG Priority Program SPP2202 ‘Spatial Genome Architecture in Development and Disease’, SPP2202 (Project number 422841138).

A.P. and K.N.N. acknowledge support from the FP7 International Training Network InteGeR, ‘Integrative Gene Regulation’ (PITN-GA-2007-214902).

A.P. acknowledges support from the Deutsche Forschungsgemeinschaft (DFG; German Research Foundation) under Germany’s Excellence Strategy – EXC-2049 – 390688087, and the Medical Research Council (MRC, UK; grant MC_U120061476).

I.I.-A. was supported by a Long-Term Fellowship from the Federation of European Biochemical Societies (FEBS).

R.F.S. is a Professor at the Cancer Research Center Cologne Essen (CCCE) funded by the Ministry of Culture and Science of the State of North Rhine-Westphalia.

R.F.S acknowledges the German Ministry for Education and Research as BIFOLD – Berlin Institute for the Foundations of Learning and Data (ref. 01IS18025A and ref. 01IS18037A).

M.N. acknowledges support from NextGeneration EU M4C2 CN00000041 CUP E63C22000940007, MUR PRIN 2022 CUP E53D23001810006, MUR PRIN PNRR 2022 CUP E53D23018360001 and computer resources from INFN, CINECA, ENEA CRESCO/ENEAGRID and Ibisco at the University of Naples.

K.N.N. acknowledge support from MRC Centenary grant.

A.G.F. and S.S. acknowledge support from MRC grants MC_U120027516 and MC_UP_1605/12.

## Figure Legends

**Supp Figure 1.**
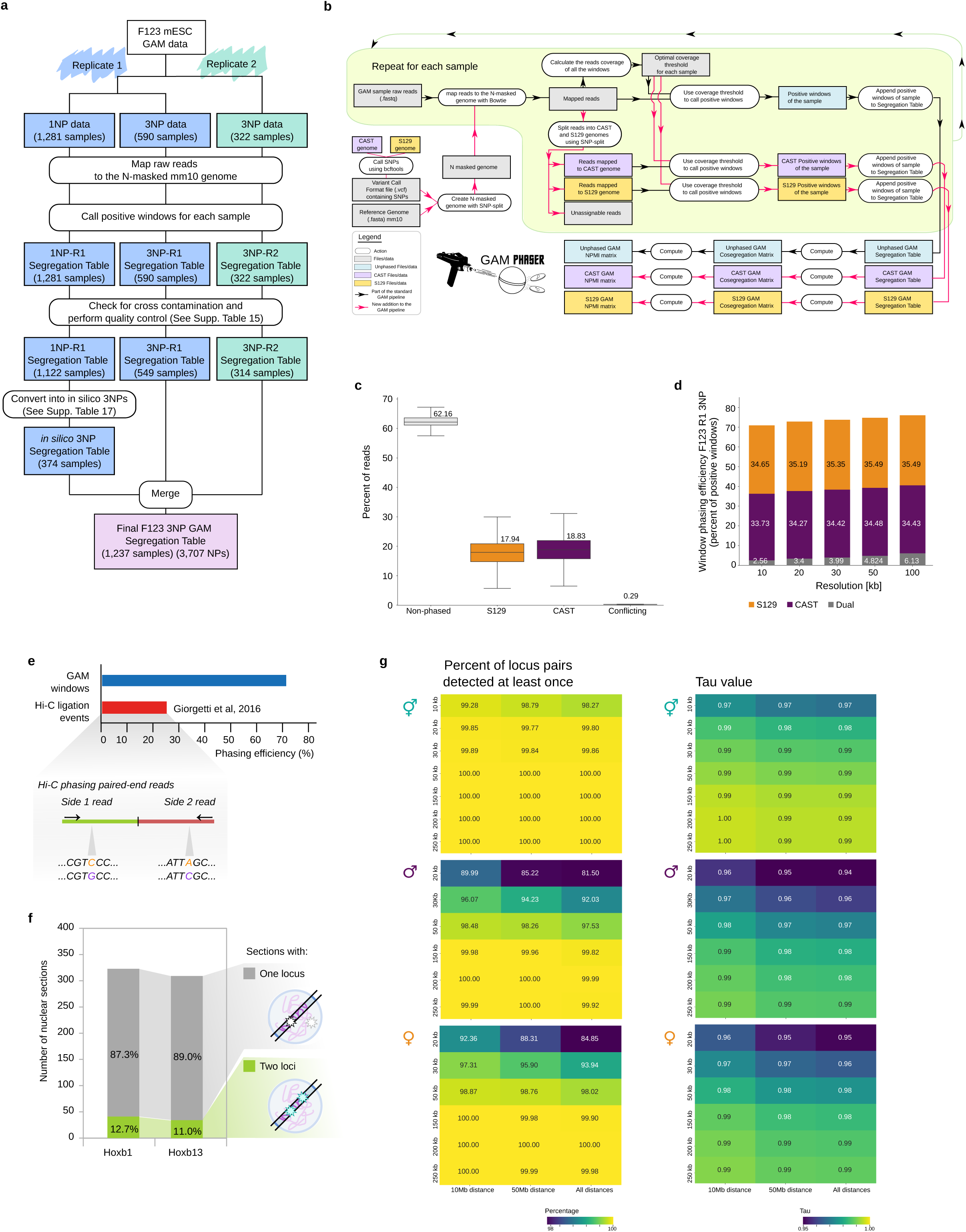
**a**, Schematic overview of GAM data collection, quality control steps, and merging of replicates R1 and R2. **b**, GAM-phaser pipeline. **c**, Percentage of reads that were phased to each allele. Conflicting reads are reads containing SNPs from both alleles. **d**, Percentage of phased positive windows in the entire segregation table for all F123 3NPs passed quality controls GAM samples. **e**, Phasing efficiency between GAM and Hi-C. GAM efficiency is measured as phased windows divided by the total number of called windows, while Hi-C efficiency is calculated dividing phased ligation events to unique ligation events; reported phasing efficiency was obtained from (Giorgetti *et al* 2016). Below, schematic of a phaseable Hi-C ligation event. **f**, Number of nuclear sections that are positive for the presence of two or one *Hoxb1* or *Hoxb13* locus detected by cryoFISH using 40 kb fosmid probes (n = 341 *Hoxb1* loci, n = 362 *Hoxb13* loci, n = 2,584 nuclear sections imaged; data source from Barbieri *et al* 2017). **g**, Percentage of locus pairs detected at least once and TAU values for different resolutions and difference distances. These metrics were used to decide on optimal resolutions of the maps.

**Supp Figure 2.**
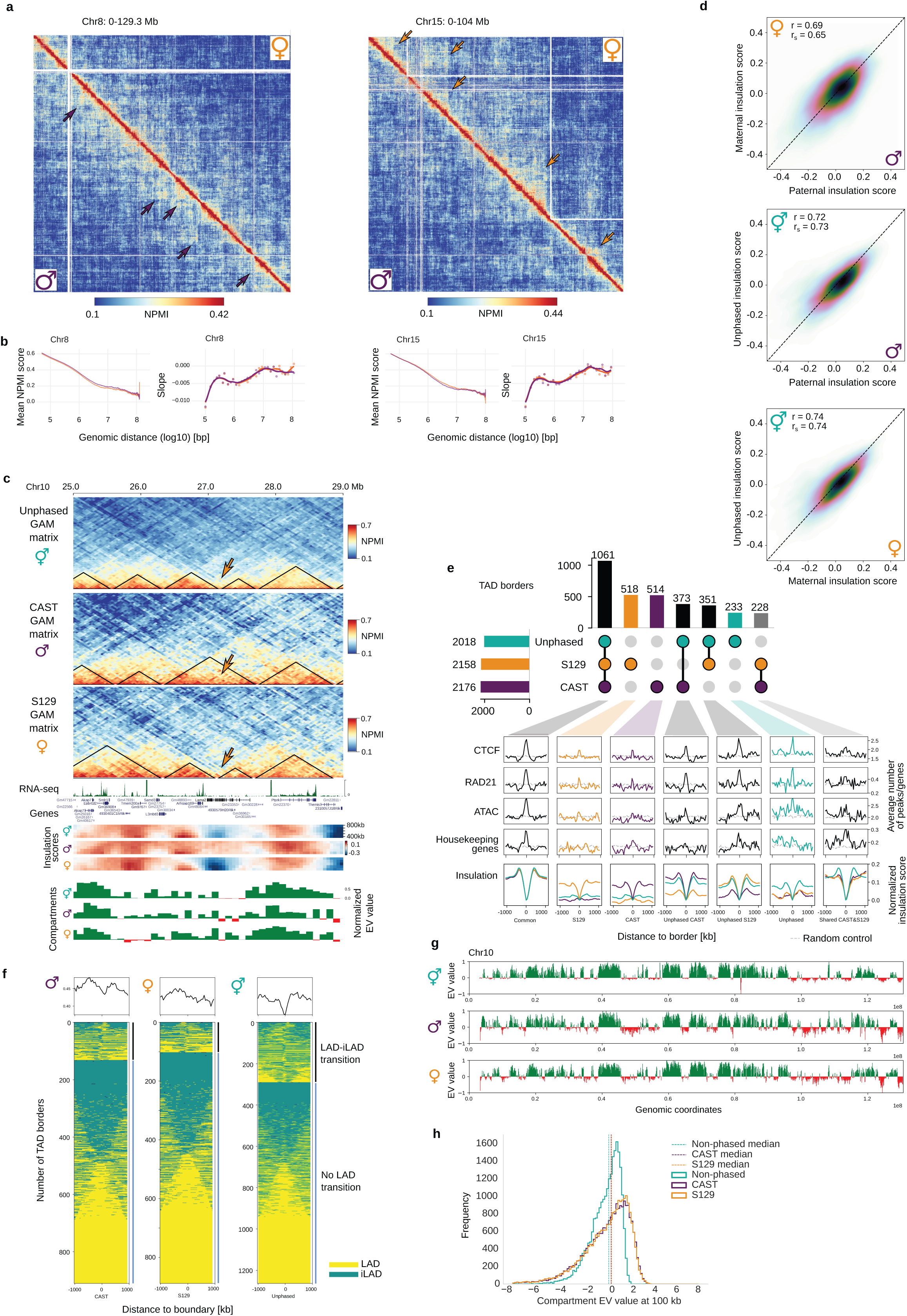
**a**, GAM matrices of chromosome 8 and 15 showing both alleles at 50Kb resolution. Colored arrows show structural differences between alleles. **b**, Distance decay curves and momentum curves for contact intensities across all distances in CAST and S129 chromosomes 8 and 15. **c**, 4 Mb region in chromosome 10 showing an allele-specific TAD border in the S129 allele. Below, RNA-seq track, insulation scores and compartment tracks for all maps. **d**, Pearson correlation coefficient (r) between combinations of CAST, S129 and unphased insulation scores at 400 kb. **e**, Upset plot of TAD border combinations between CAST, S129 and the unphased maps. Below, aggregate plots for CTCF, Rad21 and ATAC-seq peaks and housekeeping genes, centered at the TSS (± 1 kb). Normalized Insulation score is also shown for each group. **f**, Overlap of LADs and iLADs with ±1,000 kb around CAST, S129 and common TAD borders, computed from 100 kb resolution GAM matrices to match LAD annotations. Each heatmap is clustered depending on if the border overlaps with a LAD/iLAD transition or not. **g**, Compartment tracks for CAST, S129 and the unphased maps for chromosome 10. **h**, Compartments eigenvector values distribution for CAST, S129 and the unphased datasets. Discontinuous lines show the median for each dataset.

**Supp Figure 3.**
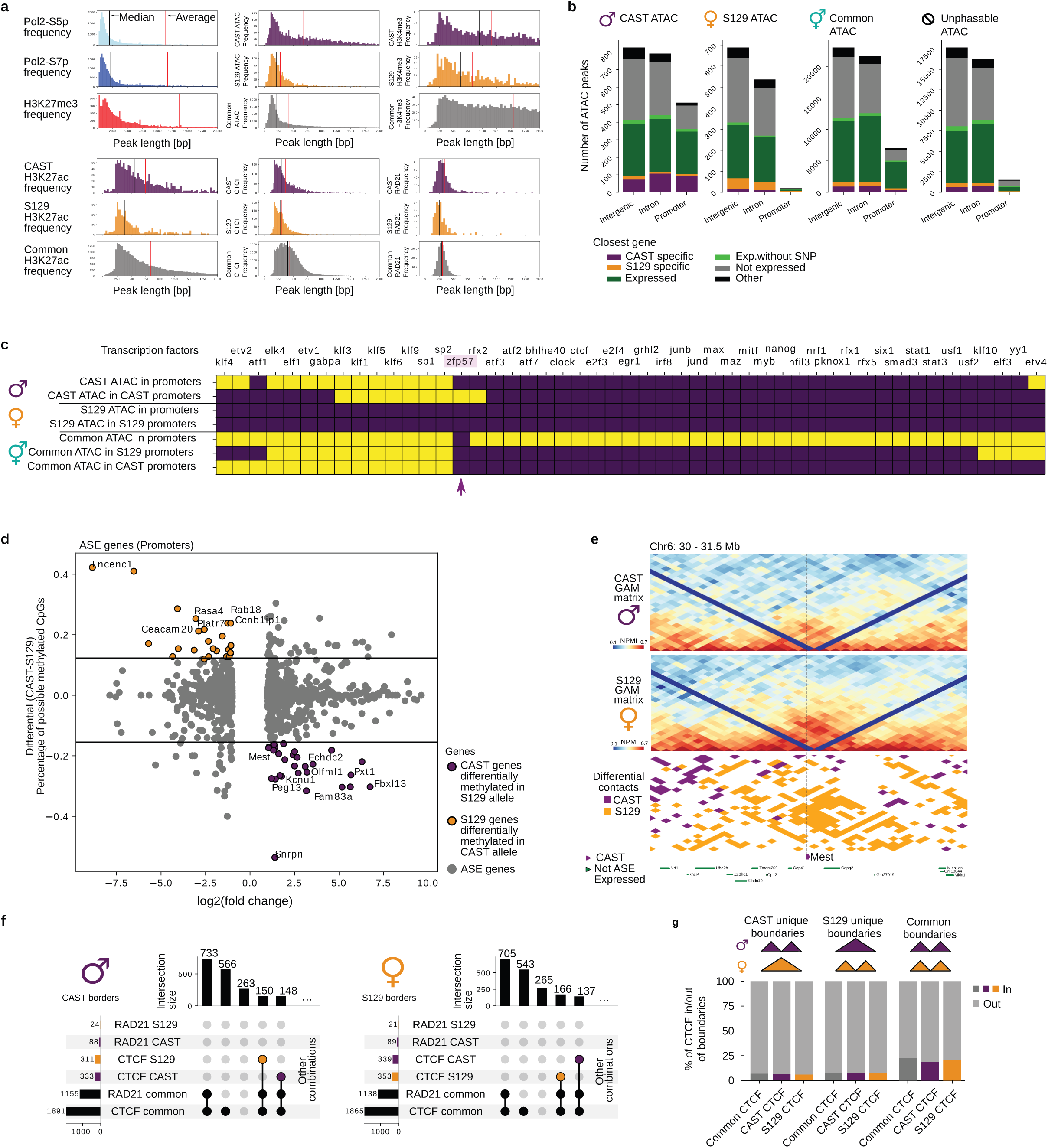
**a**, Distribution of Pol2-S5p, Pol2-S7p and H3K27me3 peaks and phased and unphased ATAC-seq, H3K4me3, CTCF, H3K27ac and Rad21 peak sizes. Black and red vertical lines correspond to the median and average of sizes for each dataset. **b**, Number of CAST, S129, common and unphasable ATAC-seq peaks in intergenic, intron and promoters. The color indicates the type (gray: not expressed; dark green: biallelic expressed; green: expressed with no SNP; orange: S129 specific; purple: CAST specific; black: unknown) of the closest gene to the ATAC peak. **c**, Heatmap showing the enriched presence (cutoffs Q-value ≤ 0.05 and p-value of ≤ 0.001) of different transcription factors that overlap with the peaks of different ATAC-seq groups. ZFP57 is the only transcription factor enriched for an allele-specific group. **d**, ASE gene promoters regarding their differential percentage of methylated CpGs. Colored are those genes with a significant amount of methylated CpGs in their promoter (top and bottom 5%) in the allele they are not expressed. **e**, CAST and S129 GAM matrices for the Mest locus (Chr6: 30-31.5 Mb). Below, differential contacts and two tracks showing CAST genes and expressed genes. **f**, Most borders contain common CTCF and RAD21 or only CTCF and each allele has a similar number of CTCF specific to either of the alleles in its borders. **g**, Percentage of CTCF peaks that are inside or outside borders.

**Supp Figure 4.**
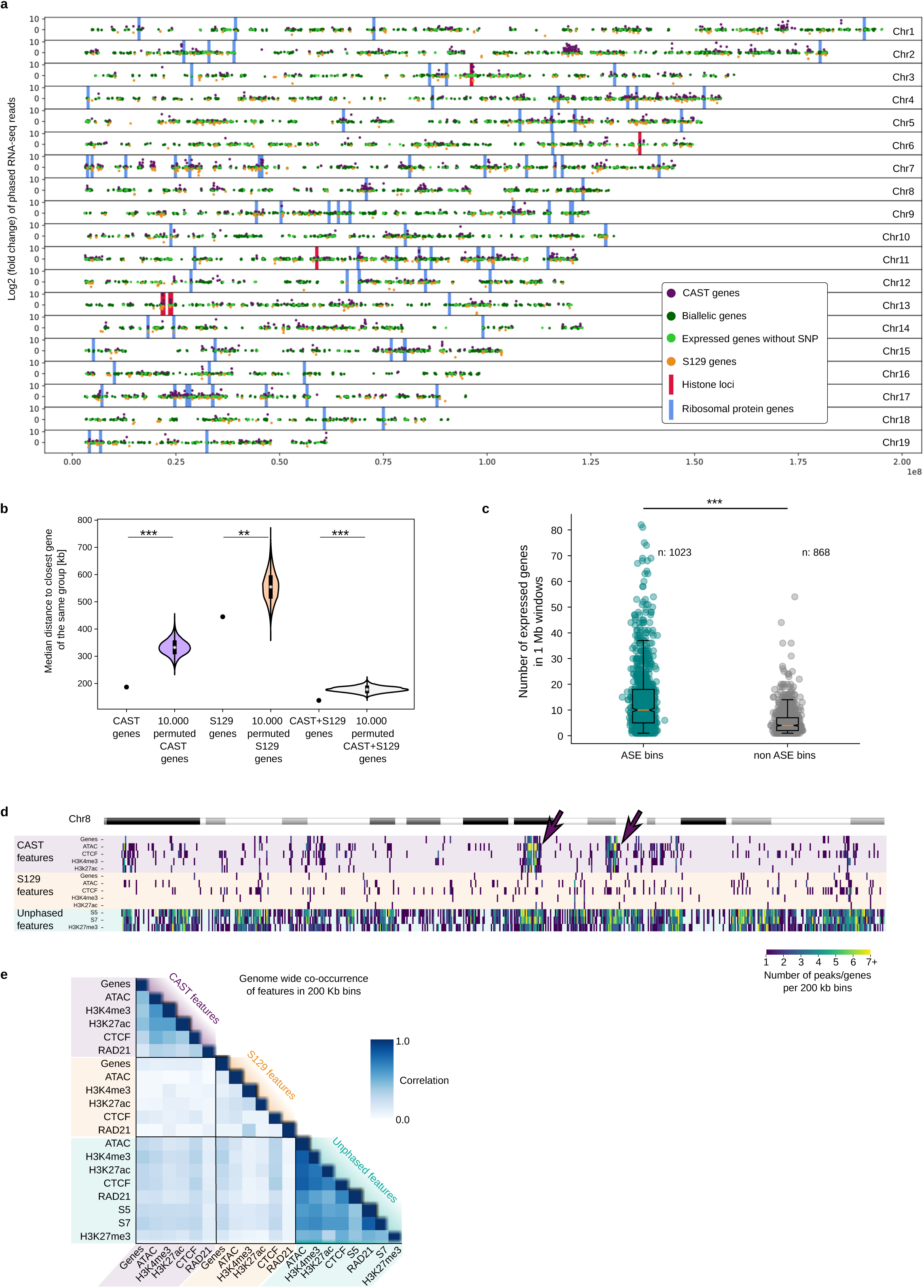
**a**, Genomic position of all expressed genes: CAST genes, S129 genes, biallelic genes and genes without SNPs. Red and Blue bars indicate the position of Histone protein genes and Ribosomal protein genes. **b**, Each of the 3 dots indicate the average distance of all genes of each type (CAST, S129 and CAST or S129) to the closest gene of that type. The violin plot shows the distribution of these averages if we permute the position of the genes 10,000 times. The permutation is carried out by randomly selecting the same number of CAST, S129 or CAST+S129 genes from all expressed genes. **c**, 1 Mb bins containing at least 1 ASE gene tend to contain more expressed genes than 1 Mb bins that do not contain ASE genes. **d**, Genomic location of allele specific features (genes, ATAC-seq, CTCF, H3K4me3 and H3K27ac peaks) and unphased features (Pol2-S5p, Pol2-S7p and H3K27me3 peaks) and their density in bins of 200kb. Arrows indicate two regions with an enrichment of CAST specific features **e**, Genome wide Pearson correlation (r) of the co-occurrence of the features in d. CAST features correlate well between each other while S129 features do not.

**Supp Figure 5.**
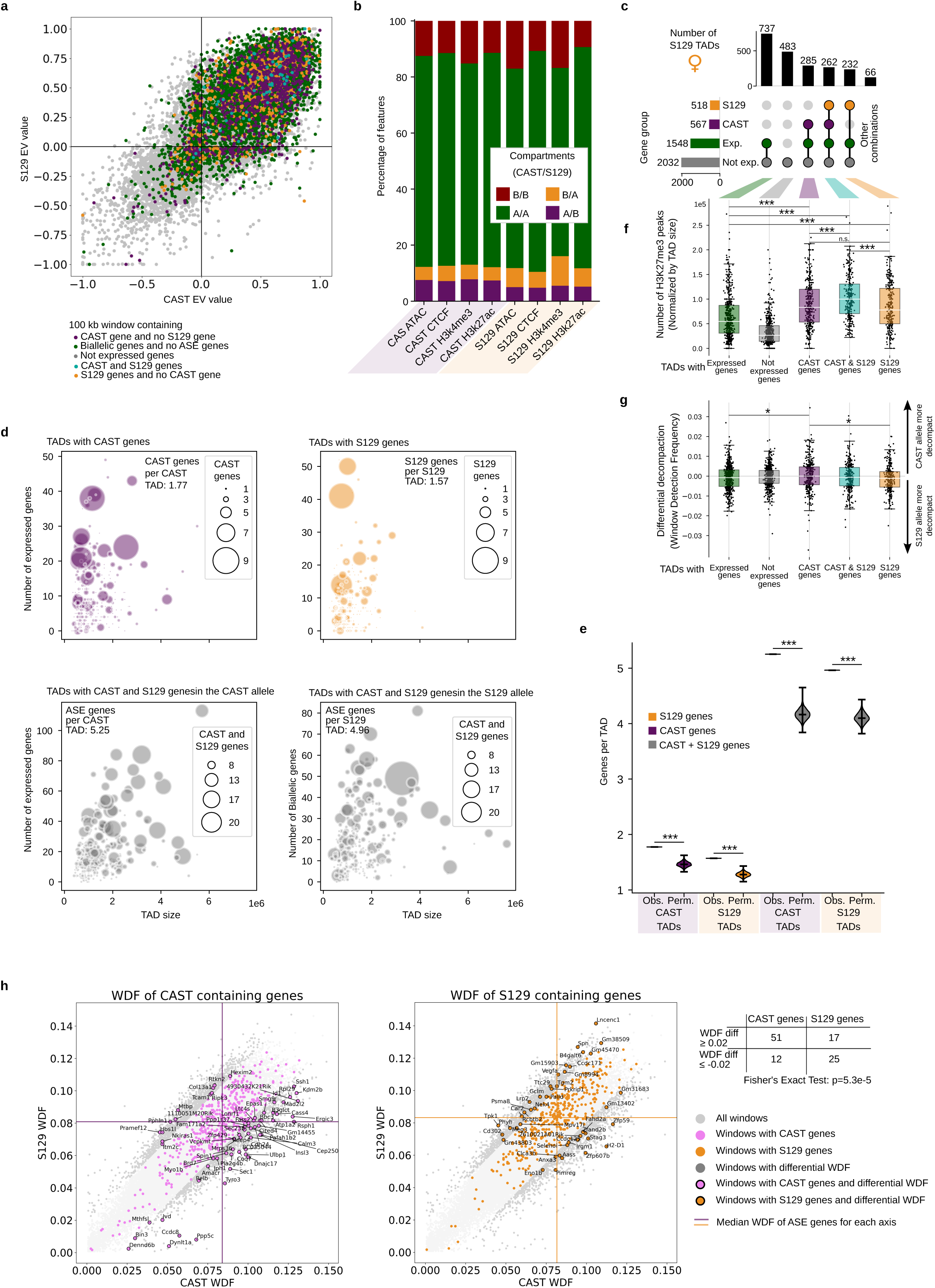
**a**, Normalized eigenvector (EV) values for the CAST and S129 allele for each 100 kb bin. Color coded, bins containing only not expressed genes, bins containing biallelic genes but not ASE genes, bins containing at least one CAST gene but not S129, bins containing at least one S129 gene but not CAST genes and bins containing at least one CAST gene and one S129 gene. **b**, Percentage of ATAC-seq, CTCF, H3K4me3 or H3K27ac peaks in each compartment combination (A and A, B and B, A and B or B and A for CAST and S129 alleles). CAST specific features show a tendency to overlap more in A/B (A specific compartment in the CAST allele), S129 specific features tend to overlap more in B/A (A specific compartment in the S129 allele.). **c**, Upset plots showing for the S129 allele, groups of TADs containing different sets of types of genes and their number. **d**, Relation between the TAD size, the number of expressed genes in a TAD, and number of genes specific to that allele (dot size) for TADs in CAST and S129. Purple refers to TADs containing CAST genes, orange to TADs containing S129 genes, and gray to TADs containing CAST and S129 genes (for both CAST allele and S129 allele, respectively). **e**, Violin plots showing the number of genes per TAD (observed, Obs.) compared to circular permutations of gene positions in the genome (permuted, Perm.). 10,000 permutations were done for each of the 4 examples in panel d and are compared to the number of genes per TAD in the original data (called *real*). All p-values are ≤0.0001. **f**, Related to panel c, number of H3K27me3 peaks normalized by TAD size (two-sided t-test: *p ≤ 0.05, **p ≤ 0.01, ***p ≤ 0.001; p-values from top to bottom in S129 TADs: 1.9×10^−14^, 3.2×10^−30^, 1.1×10^−10^, 0.00011, 1.8×10^−5^, n.s: 0.5092). **g**, Related to panel c, for each group, the differential (CAST-S129) window detection frequency. Negative values indicate decompaction in the S129 allele, while positive values indicate decompaction in the CAST allele. (two-sided t-test: *p ≤ 0.05, **p ≤ 0.01, ***p ≤ 0.001; p-values from top to bottom for S129 TADs: 0.031, 0.025. **h**, Window detection frequency (WDF) values in the CAST and S129 allele for each bin containing genes. Fisher’s exact test (p = 5.3×10^−5^) shows the significant tendency of bins with high WDF in the CAST allele containing CAST genes and bins with high WDF in the S129 allele containing S129 genes compared to bins with lower WDF.

**Supp Figure 6.**
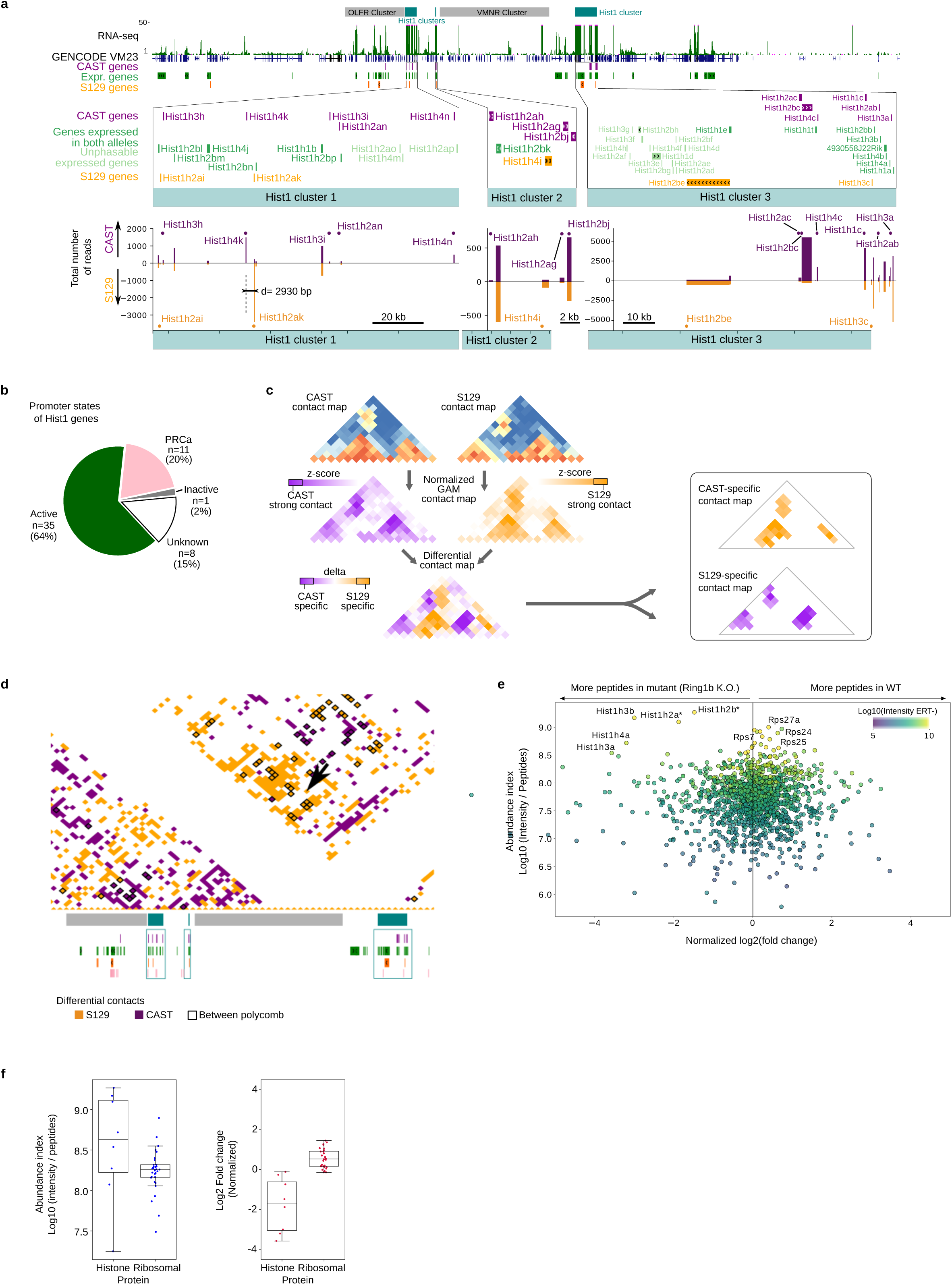
**a**, Hist1 locus region from Figure 4a with greater detail. Histone genes are depicted Number of reads phased for the CAST or S129 allele are shown for all genes in the Hist1 clusters. **b**, Proportion of Hist1 genes according to promoter state. **c**, Schematic showing the pipeline used to extract CAST and S129 specific contacts. **d**, Allele-specific contacts at the Hist1 locus as shown in Figure 4a. Black squares show the contacts where H3K27me3 peaks are present in both bins of the contact. **e**, SILAC experiments were performed in the ESC-ERT2 cells in the presence and absence of tamoxifen to induce knockout of *Ring1b* resulting in the upregulation of histone protein in the conditional knockout cells. Abundance was estimated by the ratio of intensity and number of peptides. Normalized log2 fold change was calculated applying the z score normalization to the log2 of heavy/light (H/L) ratio of the untreated experiment divided by the H/L ratio of the conditional knockout. **f**, Boxplots showing the abundance index and the log2 fold change for detected histone proteins and ribosomal proteins.

**Supp Figure 7.**
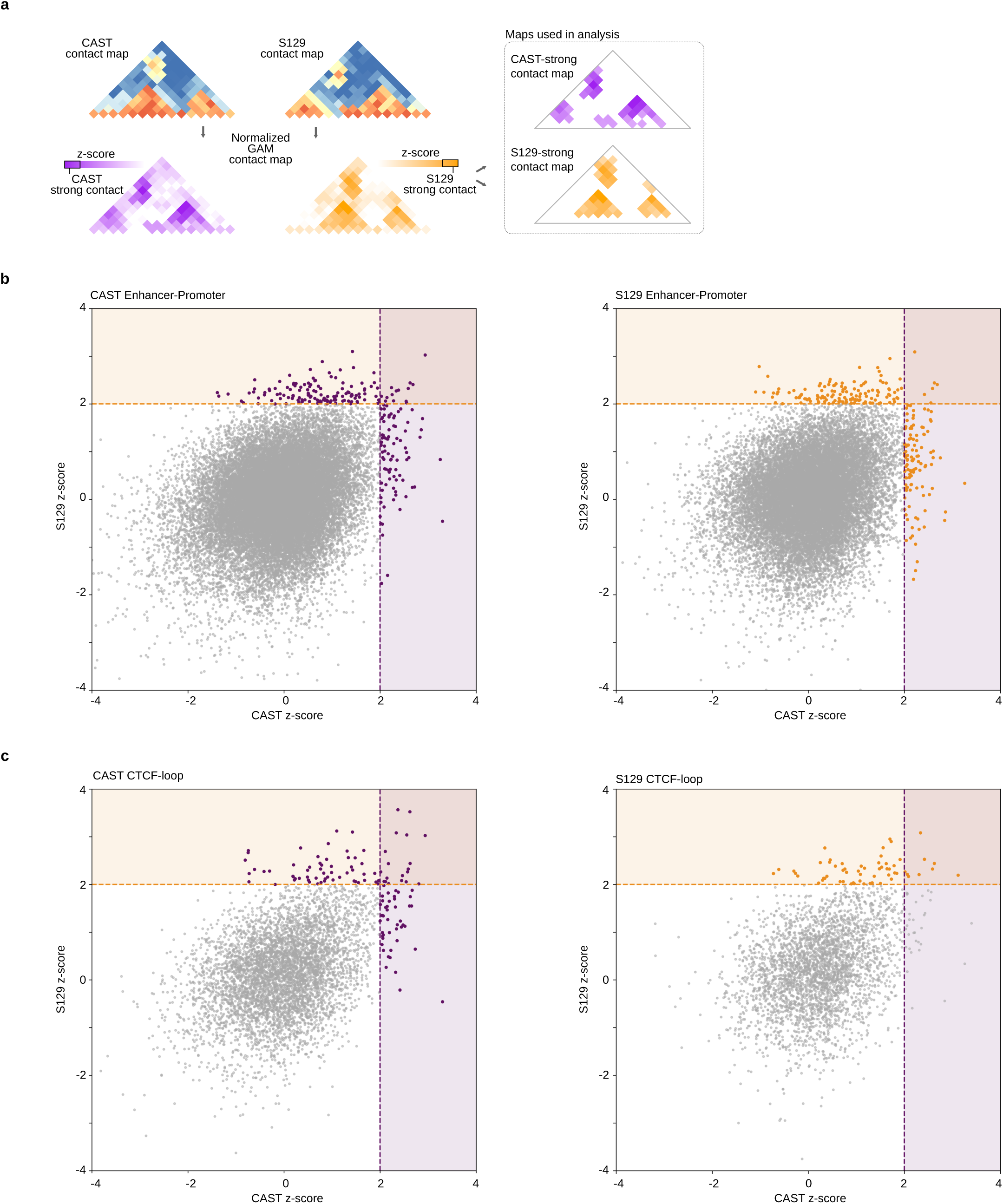
**a**, Schematic showing the strategy to identify strong allelic contacts. **b**, All possible contacts involving conditions for CAST-specific enhancer-promoter contacts and S129 specific enhancer promoter contacts. Lines mark cut-offs for strong and allele-specific contacts in each haplotype. **c**, All possible contacts involving conditions for CAST-specific CTCF loops and S129-specific CTCF loops. Lines mark cut-offs for strong and allele-specific contacts in each haplotype.

**Supp Figure 8.**
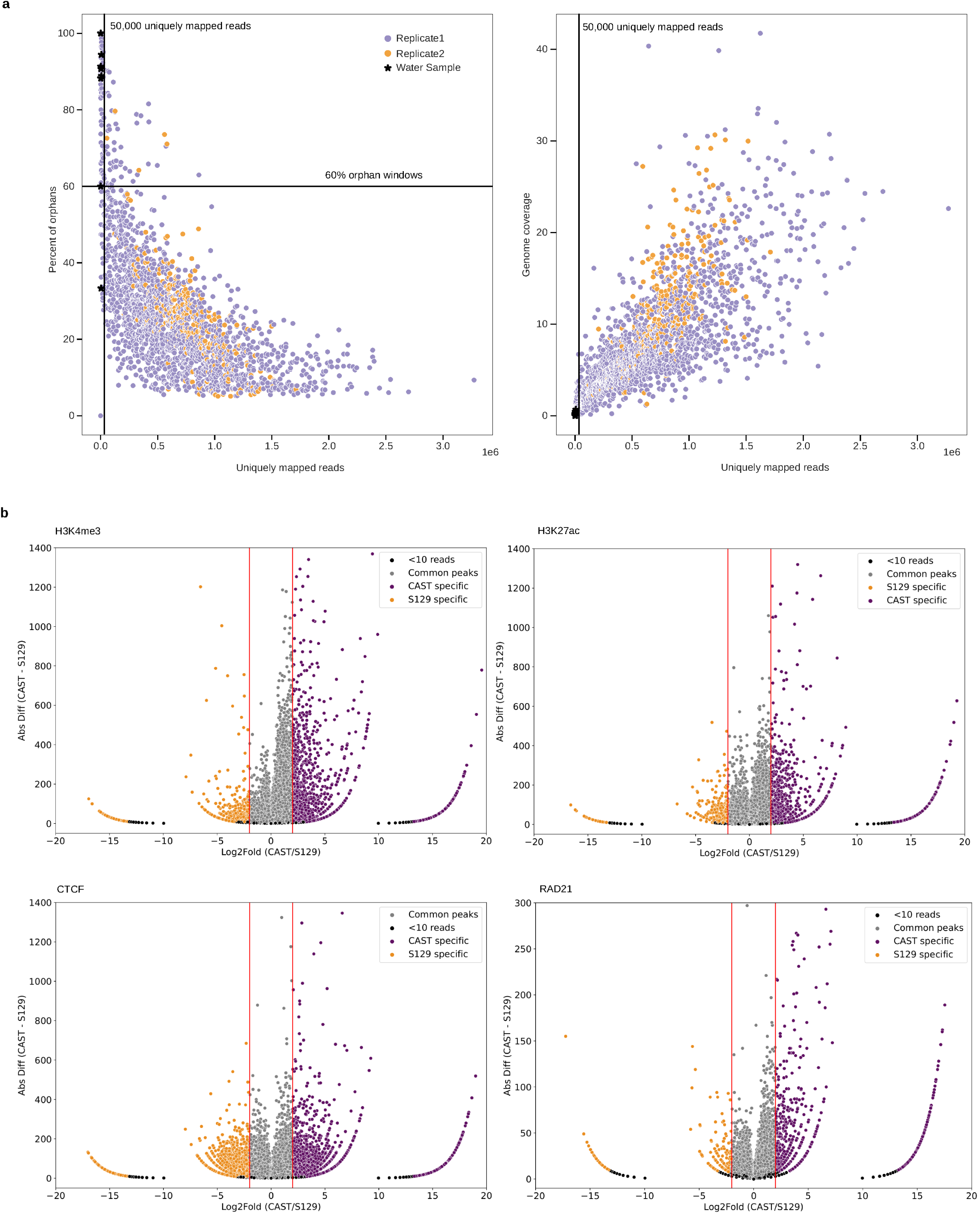
**a**, Distribution of percentage of orphan windows, uniquely mapped reads and genome coverage in each GAM sample. Replicate 1, replicate 2 and water samples are shown. Thresholds used to remove potentially bad from good GAM samples are shown in vertical and horizontal black lines **b**, Distribution of phased H3K4me3, H3K27ac, CTCF and RAD21 peaks, showing their absolute difference in phased reads (CAST – S129) and their log2 fold change.

## List of SI Tables

SI Table 1: Summary of all datasets used and produced in the study.

SI Table 2: Distances between Hoxb1 and Hoxb2 measured by cryoFISH (Barbieri et al. 2017).

SI Table 3: F123 GAM TAD borders.

SI Table 4: F123 GAM compartments and eigenvector values.

SI Table 5: Gene Ontology analysis.

SI Table 6: List of ribosomal protein genes, histone genes, housekeeping genes and imprinted genes.

SI Table 7: DNA methylated gene promoters.

SI Table 8: Genome-wide Window Detection Frequency (WDF) in the F123 GAM dataset for CAST and S129 alleles.

SI Table 9: Mass spectrometry results table.

SI Table 10: Experimental sequencing and QC metrics for GAM samples.

SI Table 11: Number of NPs produced and that passed QC.

SI Table 12: 1NP GAM sample assignment in the in-silico 3NP GAM dataset.

